# Correlated retrograde and developmental regulons implicate multiple retrograde signals as coordinators of chloroplast development in maize

**DOI:** 10.1101/2022.05.24.493305

**Authors:** Rennie Kendrick, Prakitchai Chotewutmontri, Susan Belcher, Alice Barkan

**Affiliations:** Institute of Molecular Biology, University of Oregon, Eugene, OR 97403

**Keywords:** plastid, chloroplast, retrograde signaling, maize, Arabidopsis, PTAC12, TOR

## Abstract

Signals emanating from chloroplasts influence nuclear gene expression, but roles of retrograde signals during chloroplast development are unclear. To address this gap, we analyzed transcriptomes of four non-photosynthetic maize mutants and interpreted them in the context of transcriptome dynamics during normal leaf development. We analyzed two albino mutants lacking plastid ribosomes and two chlorotic mutants with thylakoid targeting or plastid transcription defects. The ∼2700 differentially expressed genes fall into six major categories based on the polarity and mutant-specificity of the change. These distinct retrograde responses correlate with distinct developmental dynamics, with down-regulated genes expressed later in normal development and up-regulated genes acting early. Photosynthesis genes are down-regulated specifically in the albino mutants, whereas up-regulated genes are enriched for functions in chloroplast biogenesis and cytosolic translation. TOR signaling is elevated in plastid ribosome-deficient mutants and declines in concert with plastid ribosome buildup during leaf development. Our results implicate three plastid signals as integral players during photosynthetic differentiation. One signal requires plastid ribosomes and activates photosynthesis genes. A second signal reflects attainment of chloroplast maturity and represses chloroplast biogenesis genes. A third signal responds to nutrient consumption by developing chloroplasts and represses TOR, which down-regulates cell proliferation genes early in leaf development.

**One sentence summary:** Transcriptomes of non-photosynthetic maize mutants when interpreted in the context of normal developmental dynamics implicate three plastid signals as coordinators of photosynthetic differentiation.

The author responsible for distribution of materials integral to the findings presented in this article in accordance with the policy described in the Instructions for Authors (https://academic.oup.com/plcell/pages/General-Instructions) is Alice Barkan

## INTRODUCTION

Genes encoding components of the photosynthetic apparatus are distributed between the nuclear and chloroplast genomes in plants and algae, necessitating mechanisms to coordinate gene expression between the two compartments. Furthermore, the sensitivity of photosynthetic electron transport to abiotic stress serves as a sentinel to trigger adaptive changes in nuclear gene expression (Chan et al., 2016; Kleine and Leister, 2016; Brunkard and Burch-Smith, 2018). Cross talk between the nuclear and chloroplast compartments is bidirectional: nucleus-encoded proteins participate in chloroplast gene expression and signals originating in plastids influence nuclear gene expression. The latter, denoted retrograde signals, are often divided into two categories: operational and biogenic (Pogson et al., 2008). Operational signals report the status of photosynthetic reactions, and numerous such signals have been elucidated (Chan et al., 2016; Kleine and Leister, 2016; Brunkard and Burch-Smith, 2018). Biogenic signals report the status of plastid gene expression, and are hypothesized to coordinate nuclear and plastid gene expression during the differentiation of chloroplasts from their progenitor, the proplastid. Current views of biogenic signaling center on tetrapyrroles, and invoke Mg-protoporphyrin IX as a repressor of nuclear genes involved directly in photosynthesis (Photosynthesis Associated Nuclear Genes, or PhANGs) and/or heme as a positive signal to activate PhANGs (Strand et al., 2003; Woodson et al., 2011; Terry and Smith, 2013; Chan et al., 2016; Kleine and Leister, 2016; Larkin, 2016; Wu and Bock, 2021). Proteotoxic stress from misfolded proteins in the chloroplast also impacts nuclear gene expression (Ramundo and Rochaix, 2014; Llamas et al., 2017), a phenomenon that could, in principle, contribute to both biogenic and operational signaling.

The phenomenon of retrograde signaling was first recognized in studies of albino barley and maize seedlings, which were shown to have reduced mRNA levels from several PhANGs (Mayfield and Taylor, 1984; Batschauer et al., 1986; Burgess and Taylor, 1987; Rapp and Mullet, 1991; Hess et al., 1994; Börner, 2017). These observations spurred elegant genetic screens in *Arabidopsis thaliana* that revealed a diversity of signaling pathways (reviewed in Chan et al., 2016; Kleine and Leister, 2016; Brunkard and Burch-Smith, 2018). Orthologs of the few PhANGs that had been shown to be affected in albino barley and maize are similarly affected in analogous Arabidopsis material. However, little attention has been paid to retrograde signaling in monocots since the time of the early studies, so the degree to which phenomena described in Arabidopsis occur in monocots is unclear. Furthermore, although it has been assumed that biogenic signaling plays a role during normal chloroplast development, the evidence for this is limited. The most compelling evidence comes from experiments with an Arabidopsis cell culture system, which provided evidence that a signal from developing chloroplasts activates PhANGs during light-induced greening (Dubreuil et al., 2018).

The gradient of chloroplast development along the length of the seedling leaf blade in the grasses provides an opportunity to address the role of retrograde signals during the proplastid-to-chloroplast transition during normal leaf development. Cells of the intercalary meristem at the leaf base comprise a zone of cell proliferation and harbor proplastids. Cells at increasing stages of photosynthetic differentiation are arrayed in a linear fashion from base to tip. This spatial separation of chloroplasts at successive developmental stages has been exploited to describe the morphological, physiological, and molecular changes that accompany the proplastid-to-chloroplast transition in maize, barley, wheat, and rice (Robertson and Laetsch, 1974; Dean and Leech, 1982; Baumgartner et al., 1989; Li et al., 2010; Majeran et al., 2010; Wang et al., 2014; Chotewutmontri and Barkan, 2016; Loudya et al., 2021). To capitalize on this feature to understand roles of retrograde signals during leaf development and to increase understanding of retrograde signaling in monocots, we analyzed transcriptomes of four maize mutants with well-defined lesions in chloroplast biogenesis: two albino mutants, *ppr5-1* and Zm-*murE-2,* and two chlorotic mutants, Zm-*ptac12-2,* and *tha1-1*. PPR5 is required for the accumulation of two chloroplast RNAs required for chloroplast translation (*trnG-UCC* and *rpl16*); as a consequence, *ppr5-1* mutants lack chloroplast ribosomes and plastid-encoded proteins including the plastid-encoded RNA polymerase (PEP) (Beick et al., 2008; Rojas et al., 2018), which itself is critical for the buildup of the plastid translation machinery due to its role in transcription of plastid rRNAs and tRNAs (Kanamaru et al., 2001; Suzuki et al., 2003; Williams-Carrier et al., 2014). MurE and PTAC12 are found in a complex with PEP and stimulate PEP activity (Garcia et al., 2008; Pfalz and Pfannschmidt, 2013; Williams-Carrier et al., 2014). Thus, both the Zm-*murE-2* and Zm-*ptac12-2* mutants are deficient for plastid ribosomes, although the pigment, protein, and rRNA deficiencies are more severe in the Zm-*murE* than in the Zm-*ptac12* mutant (Figure 1). THA1 is a homolog of bacterial SecA and is required for the targeting of many proteins to the thylakoid membrane (Voelker and Barkan, 1995; Voelker et al., 1997; Celedon and Cline, 2013). Consequently, *tha1* mutants maintain normal chloroplast gene expression but they are pale green, with reduced levels of photosystem I (PSI), photosystem II (PSII), and the cytochrome *b_6_f* complex. All four mutants are seedling lethal due to their photosynthetic defects. By comparing the *tha1* transcriptome to the others, we hoped to distinguish effects on the nuclear transcriptome resulting from signals that reflect photosynthetic activities from those that rely more directly on chloroplast gene expression.

**Figure 1.**
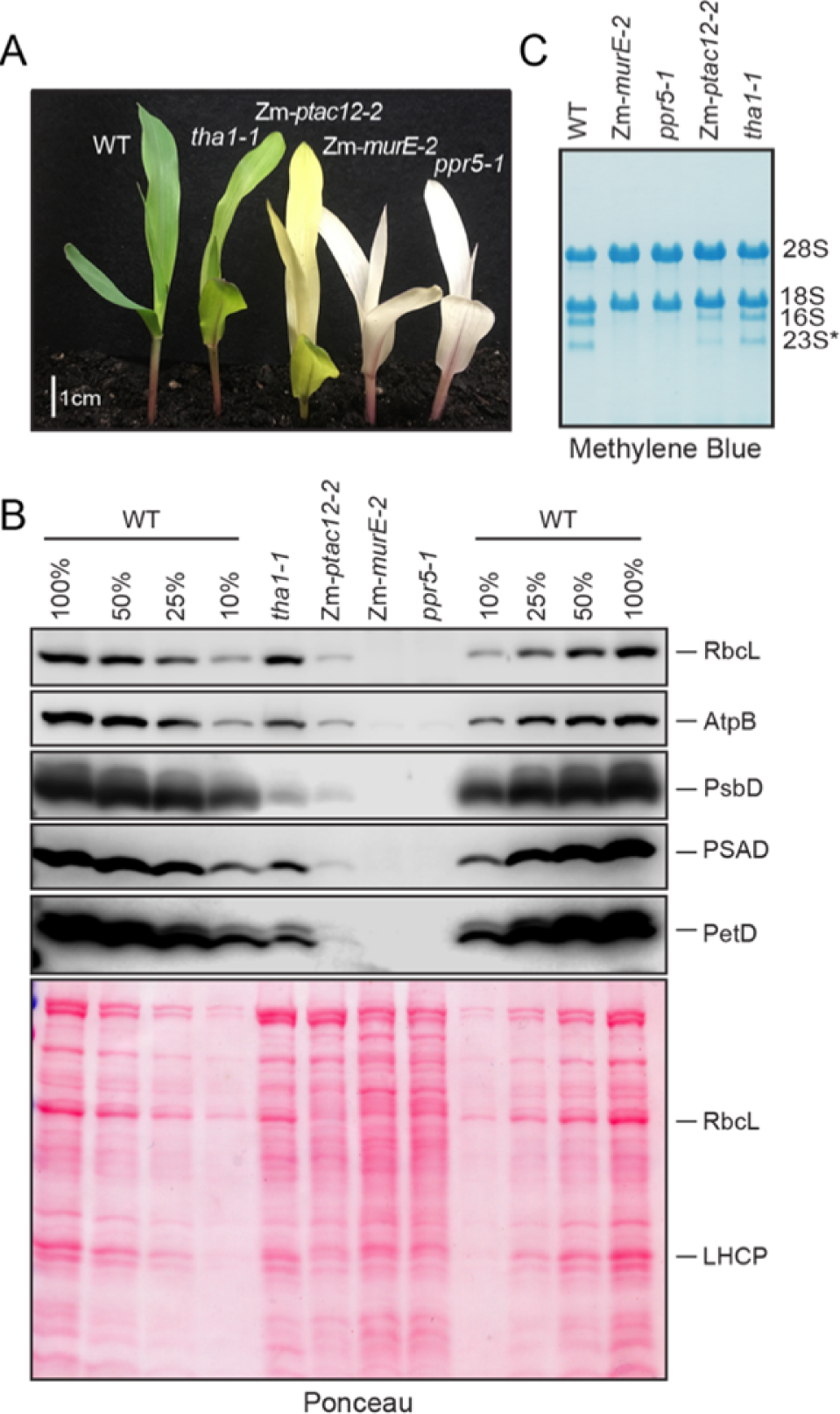
Phenotype of mutants used for RNA-seq. (A) Mutant seedlings at the developmental stage used for RNA-seq. Seedlings were grown for eight days in soil, under diurnal cycles. (B) Immunoblot analysis of components of the photosynthetic apparatus. RbcL, AtpB, PsbD, and PetD are chloroplast-encoded subunits of Rubisco, ATP synthase, PSII, and the cytochrome *b_6_f* complex, respectively. PsaD is a nucleus-encoded subunit of PSI that accumulates in proportion with plastid-encoded PSI reaction center subunits. Dilutions of extracts from two phenotypically normal siblings (WT) are shown to aid in quantification. Total proteins on the blot were imaged by staining with Ponceau S (below). (C) Abundance of chloroplast rRNAs. Total RNA was fractionated by agarose gel electrophoresis, transferred to a nylon membrane, and stained with methylene blue. Bands corresponding to cytosolic (28S and 18S) and chloroplast rRNAs (16S and 23S*) are marked. 23S* is a fragment of chloroplast 23S rRNA that arises *in vivo*.

We were also interested in using the mutant transcriptomes to clarify the role of PTAC12 in maize. The Arabidopsis PTAC12 ortholog, also known as HEMERA (HMR), localizes to both chloroplasts and the nucleus and promotes photomorphogenesis and phytochrome signaling by triggering the degradation of antagonistic Phytochrome Interacting Factors (PIFs) (Chen et al., 2010; Hernandez-Verdeja and Strand, 2018). As we have not observed morphological or molecular differences between Zm-*ptac12* mutants and maize mutants lacking other PEP-associated proteins (Williams-Carrier et al., 2014), we questioned whether the role of PTAC12 in phytochrome signaling is conserved in maize. We reasoned that such a function should be detectable as changes in the transcriptome of Zm-*ptac12* mutants that are not shared by any of the other three mutants.

Our results showed that expression of many genes encoding components of the photosynthetic apparatus (i.e. PhANGs) is strongly reduced specifically in the two mutants lacking plastid ribosomes, confirming conservation of this aspect of retrograde signaling between monocots and eudicots. However, PhANGs comprise the minority of the genes whose expression declined specifically in these two mutants. Furthermore, we observed five additional major classes of retrograde response across the mutant set, including many genes whose expression was reduced in all four mutants and many whose expression increased in all or a subset of the mutants. The up-regulated gene sets are enriched for genes involved in chloroplast gene expression, assembly of the photosynthetic apparatus, and cytosolic translation.

By comparing these varied retrograde responses to transcriptomes along the developmental gradient of the seedling leaf blade, correlations emerged between gene sets exhibiting particular expression patterns across the mutant set and regulons of normal photosynthetic differentiation. These correlations imply that several retrograde signals are integral components of the program of photosynthetic differentiation. Our findings provide evidence for two plastid-derived signals that act at distinct stages in the normal ontogeny of chloroplasts, and suggest that one signal relies on plastid translation (either directly or via its effects on PEP) and coordinates PhANG expression with that of plastid-encoded partners during the buildup of the photosynthetic apparatus, whereas the other relies on photosynthesis and represses chloroplast biogenesis genes once photosynthesis is established. Finally, we show that increased target-of-rapamycin (TOR) signaling accompanies defects in plastid ribosome biogenesis, and that TOR activity declines in concert with the activation of plastid gene expression during normal photosynthetic differentiation. These results suggest that consumption of nutritional resources by developing chloroplasts represses TOR, thereby repressing a suite of cell proliferation genes to promote exit from the cell proliferation phase of leaf development.

## RESULTS

The mutants analyzed here were described previously (Beick et al., 2008; Williams-Carrier et al., 2014; Rojas et al., 2018). They are morphologically similar to their normal siblings during their first ∼nine days of growth in soil (Figure 1A), during which growth is supported by seed reserves. The Zm-*murE-2* and *ppr5-1* mutants are albino, the *tha1-1* mutant is pale green, and the Zm-*ptac12-2* mutant is intermediate in its pigmentation. For ease of reference, both Zm-*ptac12-2* and *tha1-1* are described as “chlorotic” herein. Immunoblot analysis confirmed the expected protein deficiencies in each mutant (Figure 1B): representative stromal (RbcL) and thylakoid membrane (PetD, PsbD) proteins encoded by the chloroplast genome were not detectable in the *ppr5-1* and Zm-*murE-2* mutants and were strongly reduced in the Zm-*ptac12-2* mutant. By contrast, the *tha1-1* mutant has reduced levels of core subunits of PSI, PSII, and the cytochrome *b_6_f* complex, but near normal levels of RbcL and a membrane-extrinsic subunit of the ATP synthase (AtpB) (Figure 1B), consistent with prior results (Voelker and Barkan, 1995). The abundance of plastid rRNA was assessed by resolving leaf RNA in agarose gels and staining with methylene blue (Figure 1C). Plastid rRNA was undetectable in the two albino mutants, reduced in the Zm-*ptac12-2* mutant and at near normal levels in the *tha1-1* mutant. Taken together, these results demonstrate a profound loss of plastid ribosomes in Zm-*murE-2* and *ppr5-1* mutants, a moderate plastid ribosome deficiency in Zm-*ptac12-2* mutants, and no ribosome deficiency in *tha1-1* mutants.

RNA purified from the expanded portion of the second leaf of 8-day old seedlings (see Figure 1A) was used for RNA-seq. Each mutant was analyzed in three biological replicates. Replicate wild-type datasets were collected from phenotypically normal siblings. The RNA-seq data from replicate samples were highly correlated (Supplemental Figure S1). We used DESeq2 to quantify differential expression (Supplemental Dataset S1), and we used stringent criteria to define differentially expressed genes: an average of at least 150 reads in either the mutant or wild-type samples, a log2(fold-change) of at least 1.5, and a false discovery rate (adjusted p-value) <0.001. These criteria produced 2734 genes that were differentially expressed in at least one of the four mutants (Supplemental Dataset S2).

### Differentially expressed genes display six major types of retrograde response that correlate with distinct gene functions

We used *k-*means clustering to group genes according to their up- or down-regulation across the four mutant backgrounds. A heat map of the clustered data (Figure 2A) shows that the transcriptomes of the chlorotic mutants (*tha1* and Zm-*ptac12*) were quite similar, as were the transcriptomes of the albino mutants (*ppr5* and Zm-*murE)*, whereas the transcriptomes of the two mutant classes were readily distinguishable. However, many genes responded similarly in all four mutants, implying that much of the transcriptome response to the absence of chloroplast translation results secondarily from defects in photosynthesis. Cluster 5 had very few genes with no apparent commonalities or relationship to chloroplast processes, and will not be discussed further. The remaining clusters fell into six major groups: (i) genes that were down-regulated in all four mutants (Clusters 1, 2, 6); (ii) genes that were down-regulated primarily in the *tha1,* Zm-*ptac12* pair (Cluster 3); (iii) genes that were down-regulated specifically in the *ppr5,* Zm-*murE* pair (Cluster 4); (iv) genes that were most strongly up-regulated in the *tha1,* Zm-*ptac12* pair (Cluster 7); (v) genes that were most strongly up-regulated in the *ppr5,* Zm-*murE* pair (Cluster 8); and (vi) genes that were strongly up-regulated in all four mutants (Clusters 9 and 10). To gain insight into the functions of coregulated gene sets, we used cropPAL (Hooper et al., 2016) to catalog intracellular locations of their gene products (Figure 2B, Supplemental Dataset S3) and MapMan (Schwacke et al., 2019) to discern functional trends. A large number of MapMan bins were highly enriched in one or more cluster (Supplemental Dataset S4). Examples that capture key trends are displayed in Figure 2C. Chloroplast-localized proteins are prominent in Clusters 4 and 7, but the genes in these clusters exhibit very different retrograde responses and have distinct functional repertoires. Functions relating directly to photosynthesis are enriched only in Clusters 1 and 4, whereas other clusters represent a wide array of functions. The functional attributes of each cluster and the potential signals underlying their behaviors are discussed below, first with regard to the down-regulated gene sets followed by the up-regulated gene sets.

**Figure 2.**
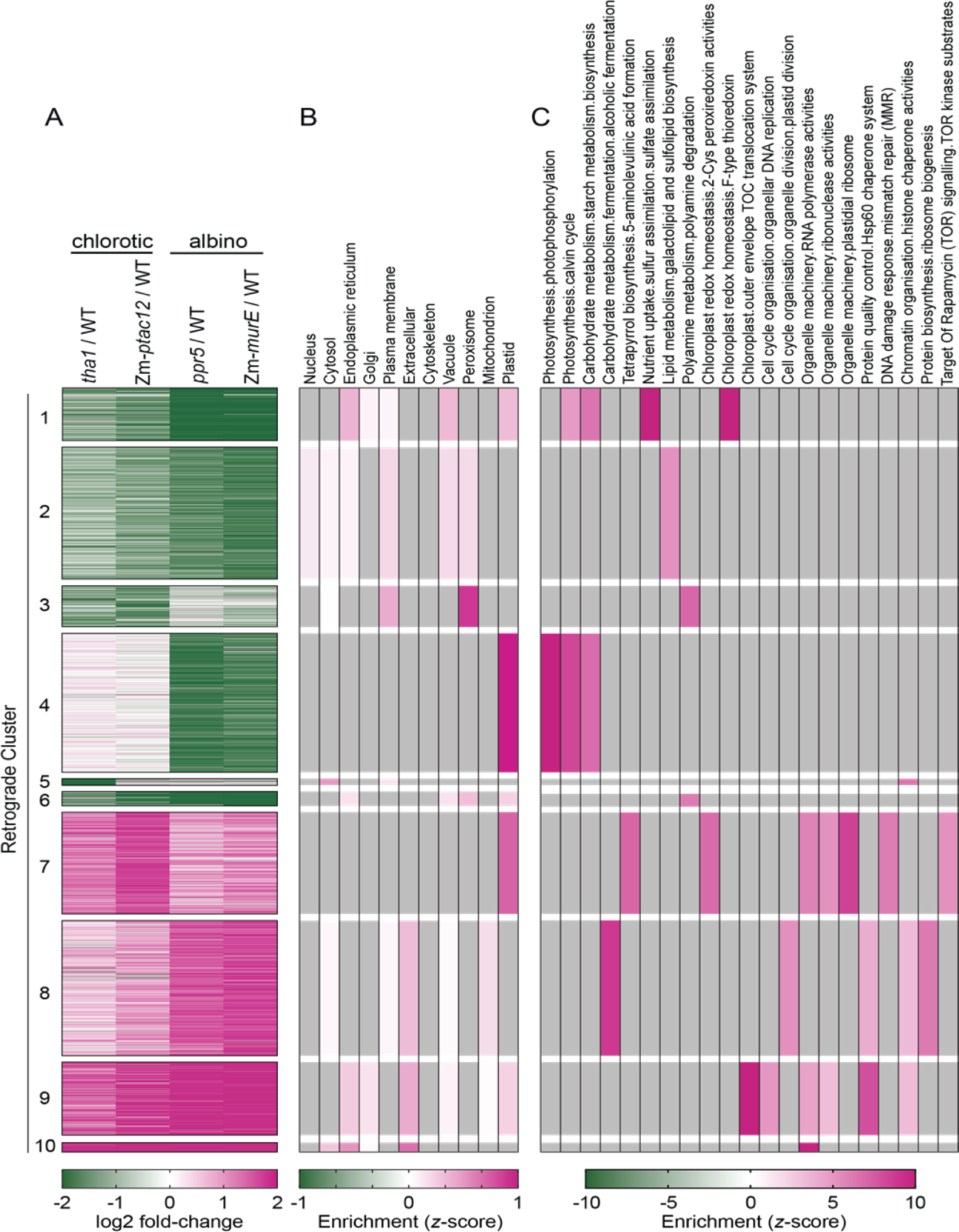
Overview of differentially expressed gene set. (A) Heat map of RNA-seq data for differentially expressed genes (log2 fold-change>1.5 and FDR<0.001 in any mutant) grouped by *k*-means clustering (*k*=10). The gap statistic supporting the use of 10 clusters is shown in Supplemental Figure S2A. (B) Heat map showing trends in intracellular locations imported from cropPal (Hooper et al., 2016). *z*-score values below zero are in gray. The complete cropPal dataset is provided in Supplemental Dataset S3. (C) Heat map displaying enrichment of selected MapMan terms. All displayed terms have a *z-*score >5 and FDR <0.01 in at least one cluster based on a simulated background distribution, but many additional terms met these criteria (Supplemental Dataset S4). Within the displayed categories, all *z*-score values > 0 with FDR<0.05 are shown. Categories that did not meet these criteria are in gray.

### Multiple signals underlie reduced nuclear gene expression in chloroplast biogenesis mutants

Cluster 4 (down-regulation specifically in the albino *ppr5,* Zm-*murE* pair) is strongly enriched for photosynthesis-related genes - particularly genes associated with photophosphorylation (Figure 2C). This cluster includes the majority of genes that are typically referred to as PhANGs: that is, genes encoding structural components of the photosynthetic apparatus. In fact, 88 of the 94 PhANGs in the differentially expressed set are found in Cluster 4, with the remaining six found in Clusters 1 or 6 (reduced expression in all four mutants) (Supplemental Dataset S5). The fact that expression of most PhANGs was reduced specifically in the mutants lacking plastid ribosomes (*ppr5* and Zm-*MurE*) is consistent with work in Arabidopsis indicating that PhANG expression is regulated by a signal that responds to chloroplast translation or PEP-mediated transcription (Woodson et al., 2013; Chan et al., 2016; Kleine and Leister, 2016). It is interesting that Zm- *ptac12* mutants, despite their substantial loss of PEP-mediated transcription (Williams-Carrier et al., 2014) and plastid ribosomes (Figure 1), show near normal expression of PhANGs and other Cluster 4 genes (Figure 2, Supplemental Dataset S2). This suggests that even a low level of PEP-mediated transcription (or plastid translation) suffices to generate the signal that activates these genes. Roughly half of the genes in Cluster 4 encode proteins that do not localize to plastids, including mitochondrial, nuclear, cytosolic, and plasma membrane proteins (Supplemental Dataset S2). Therefore, retrograde regulation of PhANGs is coordinated with that of genes affecting many extra-plastidic processes.

Many genes exhibit reduced expression in all four mutants (Figure 2A), and these encompass functional classes that are distinct from PhANGs (Figure 2C). The decrease in expression in all four mutants suggests that defects in photosynthesis drive these changes. To explore this possibility, we mined data collected for maize mutants specifically lacking either PSII, the cytochrome *b_6_f* complex, or PSI that had been collected in the context of a different project. Despite the use of different growth conditions and shallower sequencing, these data resemble those for *tha1* and Zm-*ptac12* in that genes in Clusters 1, 2, and 6 were generally repressed, whereas genes in Cluster 4 were not (Supplemental Figure S3, Supplemental Dataset S2). These similarities support the view that genes in Clusters 1, 2, and 6 respond to a signal that relies on photosynthesis. In accord with this possibility, Clusters 1 and 2 include eight FLZ/DUF581 genes (Supplementary Dataset S2), which collaborate with the energy sensor SnRK1 (Nietzsche et al., 2014; Jamsheer and Laxmi, 2015; Jamsheer et al., 2018; Li et al., 2021a).

The expression of genes in Cluster 3 was most strongly reduced in the Zm-*ptac12* and *tha1* mutants (Figure 2A). This reduction correlates with the presence of chlorophyll and is shared by the Zm-*hcf136* mutant lacking PSII (Supplemental Figure S3), suggesting involvement of a signal reflecting the redox state of the electron transport chain or chlorophyll-mediated production of reactive oxygen species. A role for oxidative stress in the Cluster 3 response is supported by the enrichment of genes involved in polyamine degradation, proteins that localize to the peroxisome (the site of polyamine catabolism) (Figure 2), and the MapMan terms Reactive Oxygen Generation and Jasmonic Acid Synthesis (Supplemental Dataset S4). Together, these results suggest that chlorophyll-mediated oxidative stress contributes to the Cluster 3 response.

In summary, the varied responses of genes whose expression was reduced in our mutant set suggest that a minimum of three types of retrograde signal underlie these behaviors: one signal relies on chloroplast translation and/or PEP-mediated transcription (Cluster 4), the second reflects photosynthesis/energy status (Clusters 1, 2, 6), and the third may be modulated by oxidative stress (Cluster 3).

### Signals underlying increased expression of nuclear genes in chloroplast biogenesis mutants

Studies of biogenic retrograde signaling have emphasized genes whose expression decreases when chloroplast biogenesis is compromised. In that context, the large number of genes expressed at elevated levels in the mutants attracted our attention. Most such genes were activated in all four mutants, but they fell into several clusters with distinct retrograde responses. Cluster 7 (strongest increase in the *tha1, ptac12* pair) is particularly notable for enrichment of terms relating to chloroplast gene expression, a theme that is discussed below. Other enriched terms include chloroplast redox homeostasis and cadaverine synthesis (Figure 2, Supplemental Dataset S4), consistent with involvement of a signal related to redox state of the electron transport chain or chlorophyll-mediated reactive oxygen species. In accord with the latter possibility, Cluster 7 includes two paralogs of the singlet oxygen signaling gene *Executor 2* (Lee et al., 2007) (Supplemental Dataset S2). Terms relating to sugar metabolism and “TOR kinase substrates” are also enriched in Cluster 7, suggesting a role for energy signaling. Cluster 9 (strongly up-regulated in all four mutants) also bears signatures of energy signaling, as it includes the gamma subunit of SnRK1 (KING1) and two members of the trehalose-6-phosphate synthase family, which regulates SnRK1 activity (Li et al., 2021a). Finally, a signal resulting from disrupted protein homeostasis is suggested by the presence of the *HSP21* gene in Cluster 9, as HSP21 is a marker of a chloroplast unfolded protein response (Llamas et al., 2017). Many genes in Clusters 7, 9, and 10 are expressed at elevated levels in mutants specifically lacking PSI, the cytochrome *b_6_f* complex, or PSII (Supplemental Figure S3, Supplemental Dataset S2), supporting a role for an energy deficit in these responses.

By contrast, genes in Cluster 8, which are most strongly up-regulated in the albino *ppr5,* Zm-*murE* mutant pair, are minimally affected in mutants lacking individual photosynthetic complexes (Supplemental Figure S3, Supplemental Dataset S2), indicating that this response does not result from a defect in photosynthesis. Furthermore, Cluster 8 is distinctive with regard to the enrichment of terms related to cell proliferation: chromatin organization, cell wall organization, cytokinesis, nuclear DNA replication, cytosolic protein biosynthesis, cytosolic ribosome biogenesis, and plastid division (Figure 2C, Supplemental Dataset S4). The target-of-rapamycin (TOR) pathway is known to activate genes of this nature (Scarpin et al., 2020), suggesting that TOR signaling underlies their up-regulation in the albino mutants. This possibility is supported by experiments described below.

### Many chloroplast biogenesis genes are up-regulated in chloroplast biogenesis mutants

It is notable that all up-regulated clusters share an enrichment of MapMan terms relating to chloroplast biogenesis (Figure 2C): organelle DNA replication and the TOC plastid import system (Cluster 9), plastid division (Cluster 8), organelle RNA polymerase activities (Clusters 7, 9, 10), organelle ribonuclease activities (Clusters 7 and 9), and plastid ribosome (Cluster 7). Additional strongly enriched MapMan terms relating to chloroplast biogenesis (Supplemental Dataset S4) include organelle RNA stability, organelle RNA splicing, organelle RNA editing (Clusters 7 and 9), and the thylakoid membrane Tat translocation system (Cluster 7).

To highlight the distinct retrograde regulation of genes involved in chloroplast biogenesis and those that function directly in photosynthesis (ie PhANGs), we extracted genes with established functions in either category from the differentially expressed gene set (Supplemental Dataset S5) and displayed the data in heat maps (Figure 3A). Whereas expression of all PhANGs was strongly reduced specifically in the two mutants lacking plastid ribosomes, almost all of the genes involved in chloroplast biogenesis (import, division, gene expression, protein targeting, assembly) were expressed at elevated levels in most of the mutants. For example, of the 105 genes in our differentially-expressed set encoding proteins that function in chloroplast translation or RNA metabolism (e.g. ribosomal proteins, tRNA synthetases, PPR proteins), all but eight are up-regulated in our mutant set: 73 are found in Cluster 7 and 24 are found in other up-regulated clusters (8, 9, 10) (Supplemental Dataset S5). Genes that did not follow the typical pattern share similarities. For example, the four genes involved in plastid transcription that are coregulated with PhANGs encode sigma factors that play specialized roles in mature chloroplasts (Chi et al., 2015), contrasting with genes encoding PEP-associated proteins (e.g. PTAC3, PTAC10, FLN1, FLN2), which are expressed at elevated levels in all mutants (Figure 3A). In the plastid translation and RNA metabolism group, the few characterized genes that are coregulated with PhANGs stand apart in the developmental timing of their activity: these encode CSP41A and CSP41B, whose functions are limited to mature chloroplasts (Leister, 2014), and TPJ1, which activates synthesis of PsbJ (Williams-Carrier et al., 2019), a protein that facilitates binding of the light harvesting complex to the PSII core late during chloroplast development (Suorsa et al., 2004).

**Figure 3.**
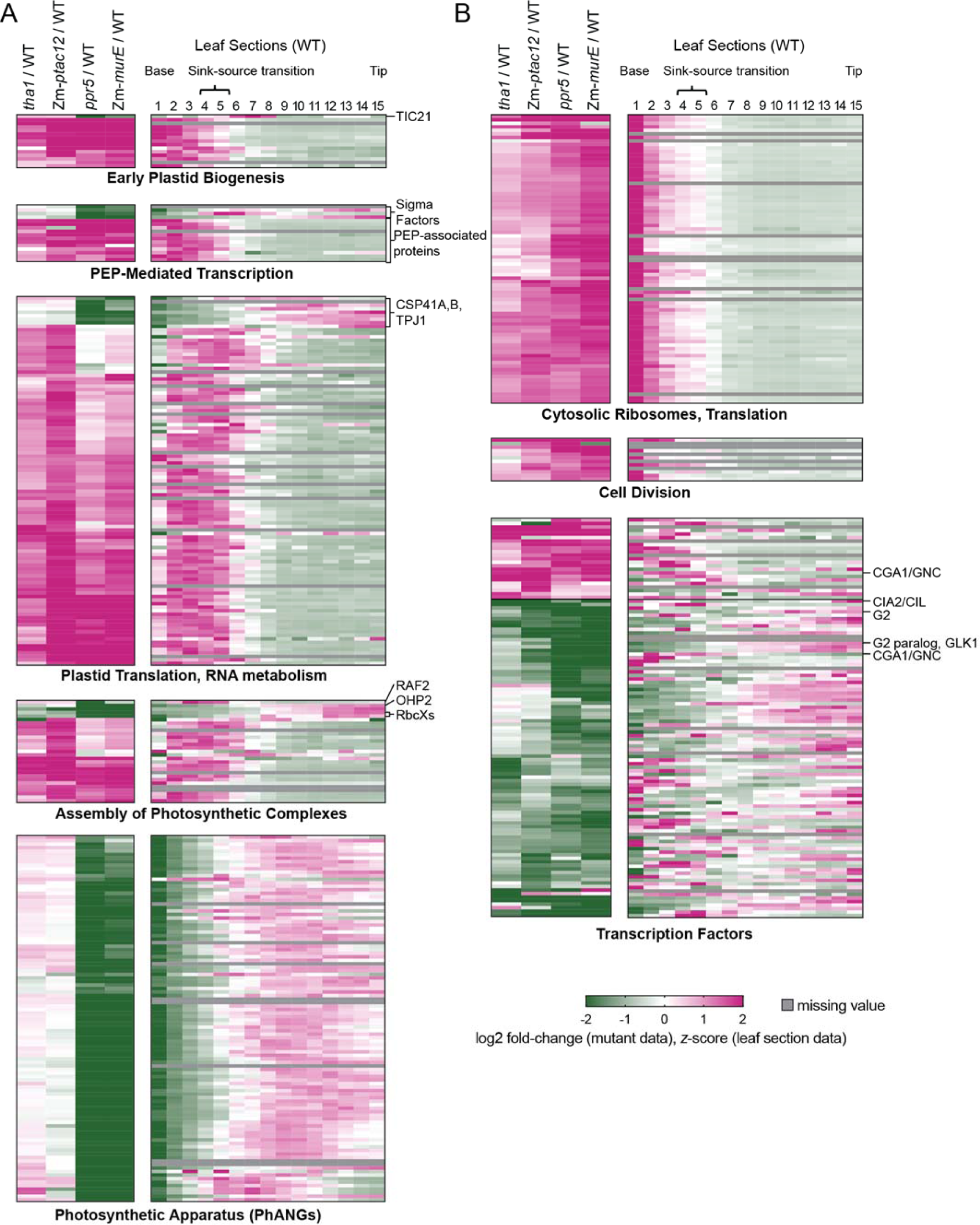
Retrograde and developmental dynamics of genes parsed by function. Heat maps illustrate differential expression in the maize mutants (left) and developmental dynamics (right) based on analysis of sections of a wild-type seedling leaf blade (Wang et al., 2014). Genes not represented in the leaf gradient data are in gray. Values and gene identifiers are provided in Supplemental Dataset S5. (A) Genes involved in chloroplast biogenesis and photosynthesis: plastid import, division, and DNA replication (Early Plastid Biogenesis), PEP-mediated transcription (sigma factors, PEP-associated proteins), Plastid Translation and RNA Metabolism (ribosomal proteins, tRNA synthetases, splicing factors, ribonucleases, PPR proteins, RRM proteins), Assembly of Photosynthetic Complexes (general chaperones, thylakoid targeting, assembly factors for individual complexes), and PhANGs. (B) Genes encoding cytosolic translation factors, cell division factors, and nuclear transcription factors.

To explore this possible relationship between the retrograde response of chloroplast biogenesis genes and their time of action during chloroplast development, the developmental dynamics of each gene are shown to the right of the mutant data in Figure 3. The developmental data were taken from a transcriptome analysis of fifteen sections along the length of a maize seedling leaf blade (Wang et al., 2014). The vast majority of the biogenesis genes peak in expression early in development (as expected), and these genes are up-regulated in the mutants. By contrast, the few genes that were down-regulated in the albino mutants (i.e. coregulated with PhANGs) peak in expression later in development (Figure 3, Supplemental Dataset S5). Examples include three proteins involved in Rubisco assembly (RAF2 and two RbcX paralogs) (Wilson and Hayer-Hartl, 2018), and the gene encoding OHP2, which is involved in PSII repair in mature chloroplasts (Li et al., 2019). The plastid import protein TIC21 appeared to be an outlier in the Early Plastid Biogenesis set, but its developmental dynamics show that TIC21 functions later in chloroplast development than typical import components.

In summary, the expression of many chloroplast biogenesis genes is altered in this mutant set. The vast majority of these are expressed at elevated levels and the few exceptions are co-regulated with PhANGs (i.e. down-regulated in the albino mutants). These distinct responses correlate with time of gene action during chloroplast development: early acting biogenesis genes are up-regulated, whereas late acting biogenesis genes are down-regulated.

### Up-regulation of genes involved in cell division and cytosolic ribosome biogenesis is associated with increased TOR activity

Cluster 8 stands apart from the other up-regulated clusters in two ways: its genes are most strongly up-regulated in the *ppr5,* Zm-*murE* mutant pair, and functions relating to chloroplast biogenesis are less prominent (Figure 2). Instead, Cluster 8 is enriched for various terms related to cell proliferation (e.g. cytosolic translation, cell division, cell wall synthesis, plastid division) and fermentation (Figure 2C, Supplemental Dataset S4). To illustrate this trend, the data for genes in the differentially expressed set with known functions in cytosolic translation, cytosolic ribosome biogenesis, nuclear DNA replication and cell division are displayed in Figure 3B. These genes are up-regulated in all four mutants, but the magnitude of the increase corresponds with the magnitude of their plastid translation defect: strongest up-regulation in *ppr5* and Zm-*murE*, less in Zm-*ptac12*, and least in *tha1*. These results suggest a role for a dose-dependent signal that represses Cluster 8 genes in response to plastid translation activity.

The up-regulation of genes involved in cell proliferation and cytosolic translation led us to consider the involvement of TOR signaling. TOR coordinates cytosolic ribosome biogenesis and cell proliferation with the availability of nutrients and energy (Wu et al., 2019; Brunkard, 2020). Inhibition of TOR in Arabidopsis reduces the abundance of mRNAs encoding proteins involved in ribosome biogenesis and cell cycle progression (Scarpin et al., 2020). The elevated expression of such genes in the mutants suggested, counterintuitively, that TOR was more active in the mutant than in the wild-type tissue. To address this possibility, we probed immunoblots with an antibody specific for eS6-240P, a phosphorylated isoform of cytosolic ribosomal protein eS6 that reports TOR activity (Dobrenel et al., 2016; Scarpin et al., 2020). TOR activity during normal leaf development was examined by immunoblot analysis of sections along the length of a wild-type seedling leaf (Figure 4A). The results showed the eS6-S240P signal to be highest in the basal sections, correlating with the expression of proliferation-related genes (Figure 3B, right) and the zone of cell proliferation (Loudya et al., 2021). Interestingly, a different phosphorylated eS6 isoform, eS6-237P, showed complementary developmental dynamics with increasing abundance commencing at the sink-source transition. The dependence of eS6-237P on TOR has not, to our knowledge, been reported. However, we found that treatment of maize and Arabidopsis seedlings with a TOR inhibitor had little effect on eS6-237P abundance (Supplemental Figure S4) suggesting that it is not dependent on TOR.

**Figure 4.**
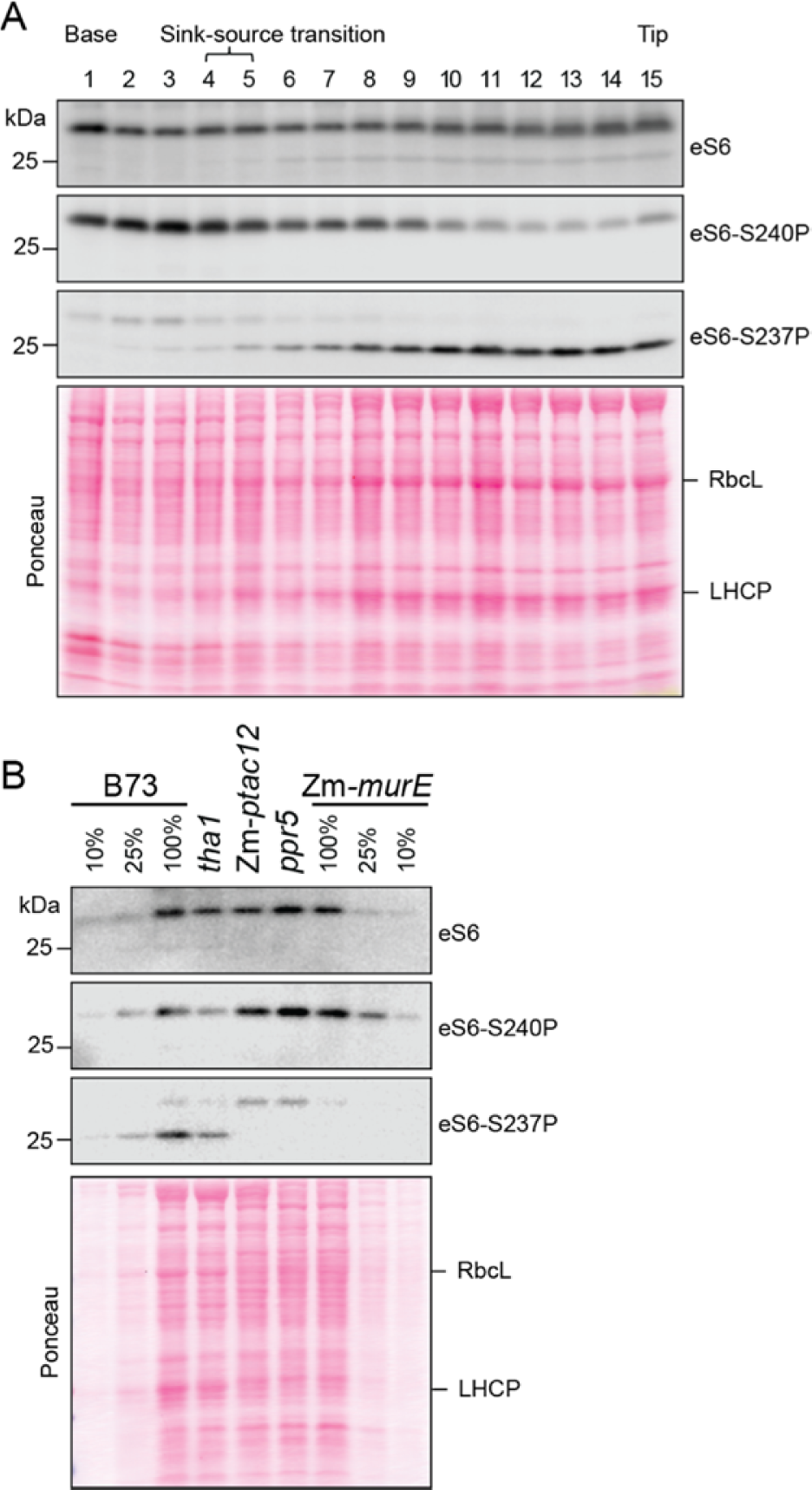
Immunoblot analysis of a marker of TOR activity during photosynthetic differentiation and in chloroplast biogenesis mutants. Immunoblots were probed with an antibody that detects cytosolic ribosomal protein S6 (eS6) (upper panels), an antibody specific for its phosphorylated S240P isoform (middle panels) and an antibody specific for its phosphorylated S237P isoform. eS6-S240P is TOR-dependent in Arabidopsis and maize (Dobrenel et al., 2016) (Supplemental Figure S4), whereas eS6-S237P appears not to be (Supplemental Figure S4). Proteins bound to the membrane were detected by staining with Ponceau S (bottom panels). The bulk of eS6 and its S240P isoform migrate at ∼28 kDa, whereas the bulk of eS6-S237P migrates at ∼25 kDa. (A) Immunoblot analysis of sections along the length of a wild-type seedling leaf blade. Plants were grown and tissue samples were taken in the same manner as used for the transcriptome data of Wang et al (2014), which are used as a point of reference in this study. Samples were harvested three hours after dawn. The third leaf was divided into fifteen segments of equal length, and an equal proportion of each sample was analyzed. The position of the sink-source transition is based on the report of Wang et al. (2014) and was not verified in these samples. Each segment was pooled from three seedlings. (B) Immunoblot analysis of mutant tissue samples grown and harvested in the same manner as those used for RNA-seq. The analyzed tissue (expanded portion of leaf 2) corresponded to sections six through fifteen in the developmental series shown in panel (A). Samples were harvested two hours after dawn on the eighth day after planting. An equal amount of protein (or the indicated dilutions) was analyzed in each lane.

Our RNA-seq data came from the expanded portion of the leaf (segments 6 through 15 in the developmental series), which is apical to the peaks of eS6-S40P (Figure 4A) and cell proliferation gene expression (Figure 3B) in wild-type tissue. Immunoblot analysis of tissue analogous to that used for RNA-seq (Figure 4B) showed the eS6-S240P signal to be stronger in the Zm-*ptac12, ppr5,* and Zm-*murE* samples than in the wild-type or *tha1*, whereas the eS6-237P signal was weaker. Thus, the *ppr5,* Zm-*murE*, and Zm-*ptac12* data, although coming from the apical region of the leaf, mimic that near the leaf base in the wild-type. These findings suggest that the elevated expression of ribosome biogenesis and cell cycle genes in Cluster 8 results from maintenance of TOR activity beyond the cell proliferation zone in which it is ordinarily most active.

The elevation of TOR activity in the Zm-*ptac12, ppr5,* and Zm-*murE* mutants was initially surprising as TOR is commonly viewed as active when energy stores are replete, yet the mutants are more likely to be in an energy deficit. However, TOR integrates multiple cues, and its activity is positively correlated with concentrations of free amino acids and nucleotides (Brunkard, 2020). The biogenesis of chloroplast ribosomes and the photosynthetic apparatus consume vast nutritional resources: translation comprises less than 1% of the cellular translational output at the maize leaf base, but ramps up rapidly to ∼25% at the sink-source transition (Majeran et al., 2010; Chotewutmontri and Barkan, 2016). Mutations or treatments that block the production of plastid ribosomes would prevent the nutrient depletion that likely accompanies this early chloroplast biogenesis phase, resulting in high TOR activity beyond the basal region in which it is typically most active. Indeed, the up-regulation of cell proliferation and other genes in Cluster 8 is, in general, inversely correlated with the abundance of chloroplast ribosomes: Cluster 8 genes are minimally affected in *tha1*, more so in Zm-*ptac12*, and most strongly in the two albino mutants (Figure 2A). This, together with the inverse correlation between TOR activity and plastid ribosome buildup early in normal chloroplast development (Figure 4A), suggest that the consumption of amino acids and nucleotides during the build-up of the chloroplast gene expression machinery represses TOR, providing a link between chloroplast biogenesis and the transition from cell proliferation to expansion during leaf development.

### Retrograde regulons correlate with developmental dynamics during photosynthetic differentiation

Transcriptome and proteome analyses of the developmental gradient along the seedling leaf blade in cereals (Majeran et al., 2010; Wang et al., 2014; Loudya et al., 2021) provided evidence that the differentiation of proplastids into chloroplasts is programmed by successive waves of expression of genes involved in different steps. Genes involved in the maintenance and proliferation of proplastids peak in expression in the most basal leaf sections. These decrease in expression a short distance above the leaf base, coinciding with the activation of genes involved in the buildup of the chloroplast genetic system and assembly of the photosynthetic apparatus. Expression of chloroplast biogenesis genes then declines toward the leaf tip, in concert with the activation of genes involved directly in photosynthesis (Loudya et al., 2021). Results described above suggested correlations between these dynamics of gene expression during normal photosynthetic differentiation (“developmental regulons”) and particular patterns of gene expression across the mutant set (“retrograde regulons”). For example, most chloroplast biogenesis genes are up-regulated in the mutants, with the few exceptions mimicking both the retrograde and developmental regulation of PhANGs (Figure 3A).

To comprehensively examine relationships between retrograde and developmental regulons, we compared the retrograde response of every gene in our differentially expressed set to its developmental dynamics along the wild-type leaf blade (Figure 5A). A visual comparison of the two datasets suggests that the majority of genes exhibiting similar retrograde responses indeed display similar developmental dynamics. To quantify these correlations, we used *k-*means clustering to group genes according to their developmental dynamics (Supplemental Figure S5), annotated each gene in the differentially expressed set with the developmental cluster to which it belongs (Supplemental Dataset S2), and calculated the degree to which each developmental cluster is enriched in each retrograde cluster (Figure 5B). This quantitative approach supports the qualitative comparisons: that is, genes with reduced expression in the mutants typically increase in expression toward the wild-type leaf tip, whereas genes with increased expression in the mutants typically peak in expression early in normal leaf development (toward the leaf base). Retrograde Cluster 3 stands apart as the only cluster that did not show significant enrichment of a developmental cluster (Figure 5B). This, in conjunction with Cluster 3’s enrichment of functions related to peroxisomes, polyamine degradation, reactive oxygen generation, and jasmonic acid synthesis (Figure 2 and Supplemental Dataset S4), suggests that Cluster 3 genes respond to stress signals in a manner that is not coordinated with developmental programs.

**Figure 5.**
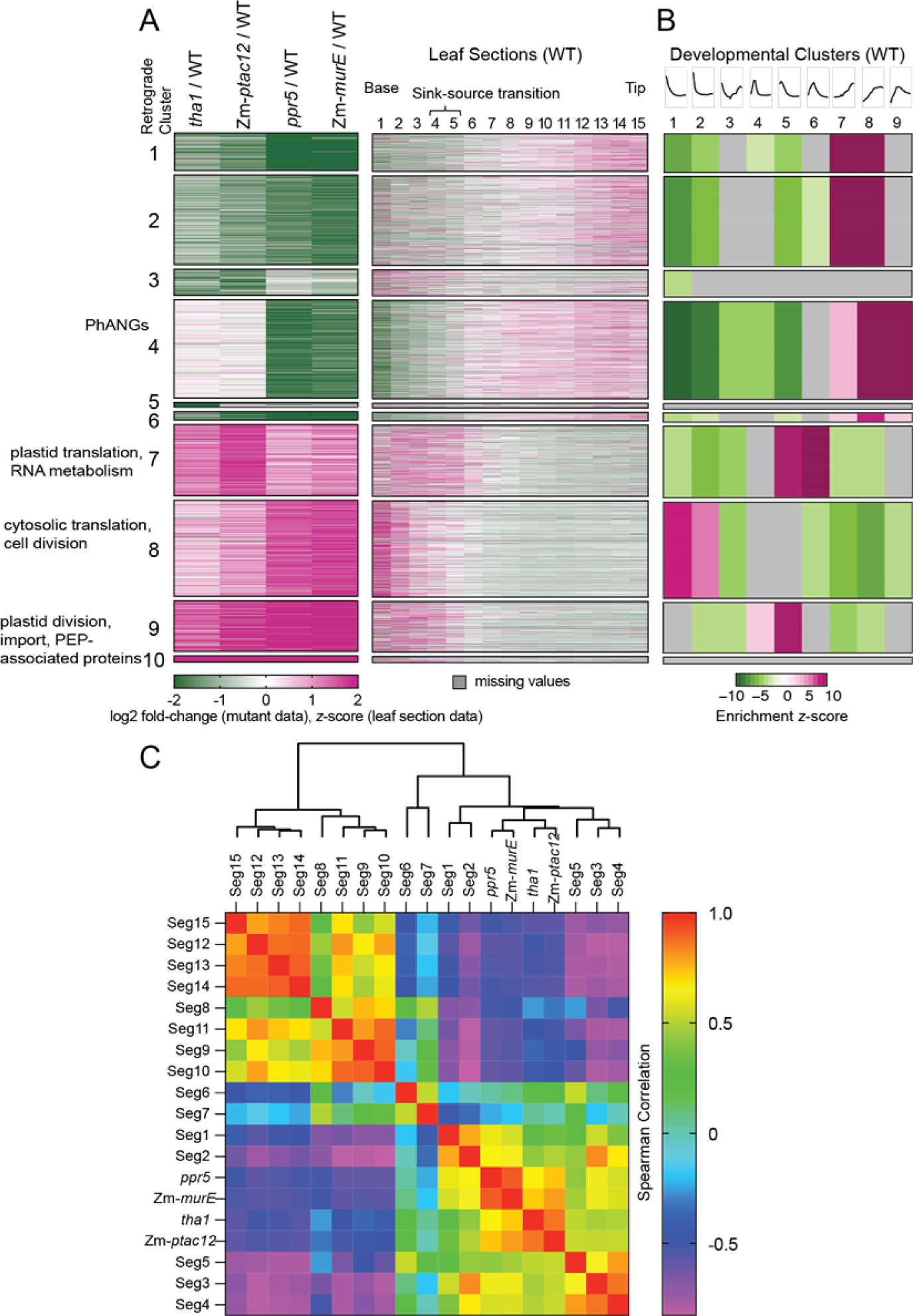
Development dynamics of genes in each retrograde cluster. (A) Heat maps showing RNA-seq data from 15 sections along the length of the maize seedling leaf blade (Wang et al., 2014) (right) in the context of the retrograde clusters defined here (left). Values for genes not represented in the developmental study are in gray. Functional characteristic of clusters that are of particular relevance are indicated to the left. (B) Enrichment of developmental clusters in each retrograde cluster. Rows correspond to the retrograde clusters shown in (A). Columns correspond to developmental clusters (Supplementary Figure S5) with expression profiles diagrammed at top (left-base, right-tip). Developmental clusters that were not significantly over- or under-represented (i.e. those with FDR > 0.05) are in gray. (C) Correlation analysis of mutant transcriptomes and leaf section transcriptomes. The analysis used the differentially-expressed gene set excepting genes in Cluster 3 (see Results for rationale). The *z*-scored values for each gene in each leaf section in the developmental series were compared with the fold-change values for the same gene in each mutant transcriptome. Correlation values are provided in Supplemental Dataset S7.

Nuances of the data parse these correlations further. Thus, genes in retrograde Clusters 1 and 2 (reduced expression in all four mutants) peak in expression very near the leaf tip, whereas those in retrograde Cluster 4 (reduced expression only in mutants lacking plastid ribosomes) show lower expression in the base and a more gradual increase toward the tip. Genes in retrograde Clusters 7 and 9 show peak expression in the basal five sections, but with the peak at a slightly earlier stage in retrograde Cluster 9 than in Cluster 7 (Figure 5A and 5B). These dynamics correlate with functional enrichment of genes involved in different stages of chloroplast biogenesis - either early plastid biogenesis (e.g. import and division in Cluster 9) or chloroplast translation and photosystem assembly (Cluster 7). Genes in retrograde Cluster 8, with highly elevated expression primarily in the mutants lacking plastid ribosomes, show peak expression at the very base of the leaf where cells are proliferating, consistent with the enrichment of genes involved in cell division, DNA replication, and cytosolic ribosome biogenesis. The coherence of the relationships between retrograde and developmental regulons instills confidence in the biological relevance of the varied retrograde responses that emerged from our data.

The down-regulation of genes that act late during normal chloroplast development and up-regulation of genes that act early indicate that chloroplast biogenesis mutants maintain transcriptomes that are characteristic of early stages of photosynthetic differentiation. This is in accord with a hypothesis that was put forward based on analysis of a handful of molecular and cell biological features in the Arabidopsis *cue8* mutant (Loudya et al., 2020). To better define the stage in normal development that is mimicked in each mutant, we calculated correlation coefficients for each mutant transcriptome in comparison with each leaf section transcriptome: fold-change values of each differentially-expressed gene were compared with *z-*scores of the same gene at each stage in the developmental series (Figure 5C, Supplemental Dataset S7). The 160 genes in Cluster 3 were excluded from this analysis due to evidence that they respond to a stress signal that is not coupled to development (see above). The results for the remaining 2574 genes show that the *ppr5* and Zm-*murE* transcriptomes most closely resemble transcriptomes of leaf sections 1 through 3 at the very base of the leaf, whereas the *tha1* and Zm-*ptac12* transcriptomes most closely resemble those in leaf sections 3 through 5, spanning the sink-source transition (Figure 5C, Supplemental Dataset S7). These results suggest that two retrograde signals play integral roles in the regulatory circuits underlying photosynthetic differentiation: the absence of plastid ribosomes inhibits progression of the developmental program beyond that characteristic of the most basal regions of the leaf, whereas the absence of functional chloroplasts interferes with progression of the developmental program beyond that characteristic of the sink-source transition.

### Comparison of maize retrograde responses to those in Arabidopsis

Much of the research on chloroplast-to-nucleus retrograde signaling has involved norflurazon- or lincomycin-treated Arabidopsis seedlings. Norflurazon inhibits carotenoid synthesis, which results in photobleaching and inhibition of plastid gene expression in the presence of light. Lincomycin inhibits plastid translation, which prevents synthesis of PEP and core subunits of the photosynthetic apparatus. Plants that are treated with either molecule lack plastid ribosomes and PEP-mediated transcription, mimicking the dysfunctions in the Zm-*murE* and *ppr5* mutants. Loss of PhANG expression following such treatments has been the primary focus of this research.

PhANG expression was reduced specifically in the Zm-*murE* and *ppr5* maize mutants, consistent with the Arabidopsis data. However, hundreds of other genes exhibited reduced expression in all four maize mutants, and many genes were expressed at elevated levels in some or all of the mutants.

To determine whether the diversity of responses we observed are represented in Arabidopsis data, we compared our data to data from four genome-wide studies in Arabidopsis: a microarray analysis of lincomycin and norflurazon treated seedlings (Woodson et al., 2013), RNA-seq analysis of norflurazon treated seedlings (Zhao et al., 2019), a microarray analysis of the Arabidopsis *pap7* mutant (Grübler et al., 2017), and a microarray analysis of the Arabidopsis *psad1* mutant (Glässer et al., 2014). The *pap7* mutant is similar to Zm-*ptac12* in that both PAP7 and PTAC12 facilitate PEP-mediated transcription, and both mutations reduce but do not eliminate chloroplast translation and the accumulation of photosynthetic complexes (Figure 1) (Gao et al., 2011). The *psad1* mutant is PSI-deficient due to mutation of a gene encoding a PSI subunit, but retains some PSI activity due to a functionally-redundant paralog (Ihnatowicz et al., 2004). The *psad1* mutant resembles maize *tha1* in that it is compromised in photosynthesis. That said, the photosynthetic defect in *tha1* is more severe as indicated by the fact that *tha1* is seedling lethal, whereas *psad1* is not.

The Arabidopsis data for putative orthologs of genes in our differentially expressed set are displayed in Figure 6. Values were not reported for orthologs of many genes in our dataset (gray bars in Figure 6), but some trends are nonetheless clear. Repression of genes in Cluster 4 (including but not limited to PhANGs) is conserved in lincomycin and norflurazon treated Arabidopsis and is less prominent in the *pap7* mutant, whereas there is a trend toward up-regulation of these genes in the *psad1* mutant. This pattern parallels that for the analogous maize mutants (*ppr5/*Zm*-murE* versus Zm-*ptac12* versus *tha1,* respectively) and is in accord with the current view that PhANG expression responds to the status of plastid gene expression. Repression of genes in Cluster 1 (reduced expression in all four mutants) is suggested in the Arabidopsis data. Cluster 1 genes that respond similarly in maize and Arabidopsis include genes encoding the transcription factor GLK1 in Arabidopsis and its co-orthologs in maize, Golden2 and GLK1, which activate PhANGs (Rossini et al., 2001; Waters et al., 2009) and play a central role in retrograde regulation of PhANG expression in Arabidopsis (Leister and Kleine, 2016; Martin et al., 2016; Hernandez-Verdeja and Strand, 2018; Grübler et al., 2021). Our data show, however, that PhANGs and GLK1 are not co-regulated in response to plastid dysfunctions: PhANG expression is minimally affected in maize and Arabidopsis mutants that retain some plastid translation activity, but the expression of GLK1 co-orthologs is reduced in all of these mutants (Supplemental Dataset S5, Figure 3B). This disconnect suggests that there may be a threshold concentration of GLK1 required for gene activation, or that PhANG activation involves GLK1 in partnership with other factors. One candidate is BBX16, whose maize ortholog was strongly down-regulated specifically in the *ppr5, Zm-murE* pair (Supplemental Dataset S5) and whose Arabidopsis ortholog was recently suggested to collaborate with GLK1 to activate PhANGs (Veciana et al., 2022).

**Figure 6.**
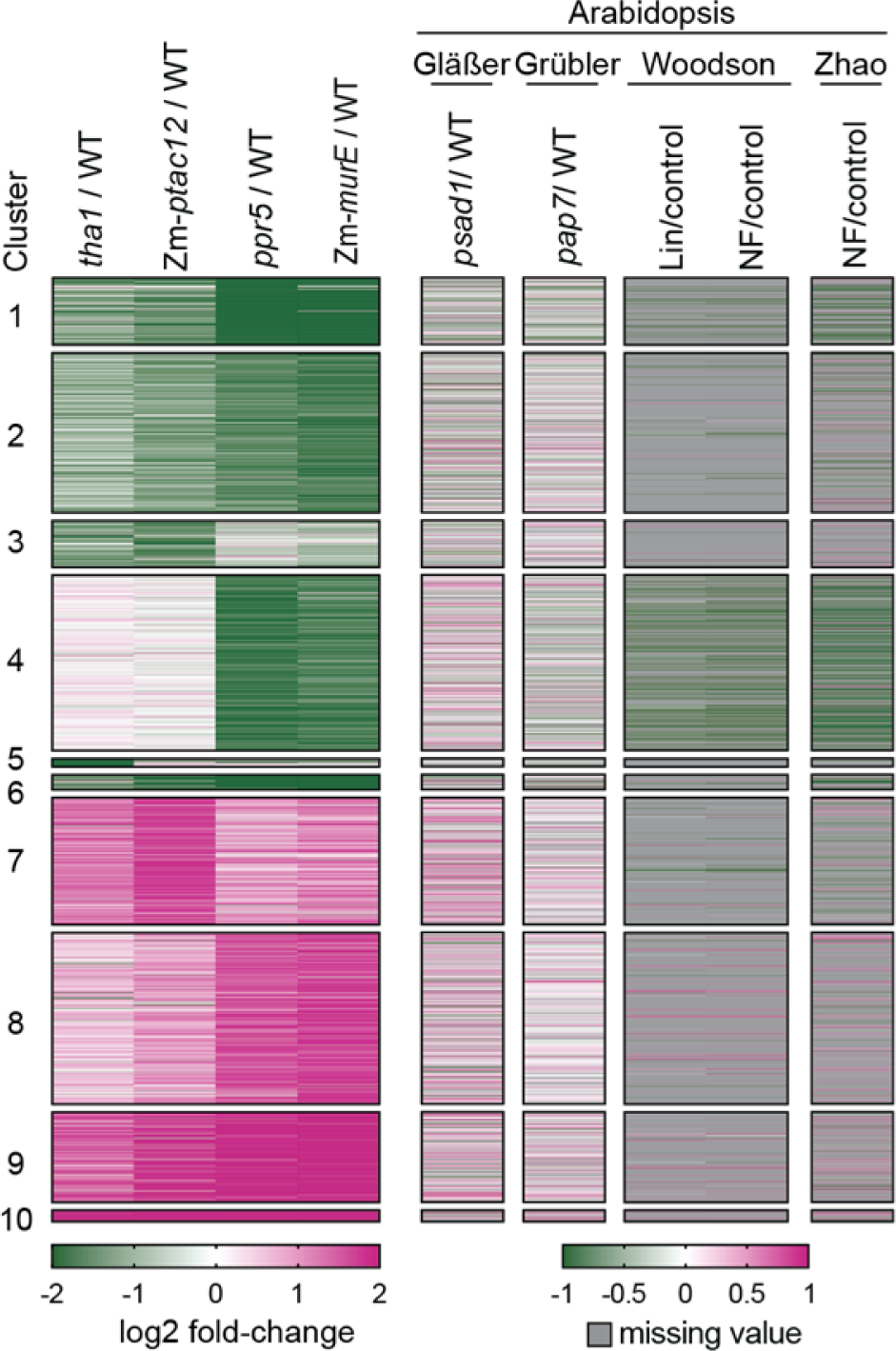
Comparison of retrograde responses in maize chloroplast biogenesis mutants to analogous Arabidopsis datasets. Data are displayed for predicted Arabidopsis orthologs of each differentially-expressed maize gene. Genes lacking a predicted Arabidopsis ortholog or for which data were not reported are in gray. The Arabidopsis data come from a microarray analysis of the *psad1* mutant (Glässer et al., 2014), a microarray analysis of the *pap7* mutant (vs wild-type in the light) (Grübler et al., 2017), a microarray analysis of lincomycin and norflurazon treated seedlings (Woodson et al., 2013), and an RNA-seq analysis of norflurazon treated seedlings (Zhao et al., 2019). The values represented in this figure are provided in Supplemental Dataset S2.

Up-regulation of some orthologous genes is also apparent in the Arabidopsis data. For example, genes in cluster 7 (most strongly up-regulated in the *tha1,* Zm-*ptac12* pair) likewise show a trend toward up-regulation in the *psad1* and *pap7* mutants but not in lincomycin- or norflurazon-treated seedlings (Figure 6, Supplemental Dataset S2). Up-regulated genes in the norflurazon and lincomycin samples include 21 genes involved in cytosolic translation, whereas such genes were not generally up-regulated in *pap7* (Supplemental Dataset S5). These results are consistent with the stronger up-regulation of cytosolic translation genes in the *ppr5, Zm-murE* mutant pair than in the *tha1, ptac12* pair.

Relatively few genes in Clusters 9 and 10 (up-regulated in all maize mutants) are shared even among the different Arabidopsis datasets. Notable exceptions include genes encoding NEP (also called RpoTp) and the PEP-associated proteins PTAC10, FLN1, and FLN2 (Supplemental Dataset S5). Genes encoding these and other PEP-associated proteins were up-regulated in maize *tha1* and Arabidopsis *psad1* (Supplemental Dataset S5), suggesting that their retrograde response results from a defect in photosynthesis and is only an indirect effect of defective plastid gene expression. Non-photosynthetic mutants in maize and Arabidopsis that maintain normal chloroplast gene expression (*tha1, pet2, hcf136* in maize, *psad1* in Arabidopsis) also exhibit elevated expression of multiple genes involved in plastid import (Supplemental Dataset S5) (Glässer et al., 2014). Together, these results provide evidence for a retrograde pathway in maize and Arabidopsis that represses genes involved in early stages chloroplast biogenesis (PEP-mediated transcription, plastid protein import) in response to a read-out of photosynthesis.

Several “core retrograde signaling modules” have been reported based on meta-analysis of transcriptomes from Arabidopsis seedlings with various chloroplast dysfunctions. One such study identified 39 differentially expressed genes shared across a variety of conditions (Glässer et al., 2014), twelve of which are represented among our differentially expressed maize genes (Supplemental Table S8). GO terms relating to the nucleolus, cytosolic ribosome, ribosome biogenesis, and plastid ribosome were enriched among genes sharing similar responses across at least three conditions (Glässer et al., 2014), suggesting conserved retrograde regulation of these functions in maize and Arabidopsis. Another study identified 152 genes whose expression changed in several conditions affecting chloroplast gene expression (Grübler et al., 2021).

Predicted orthologs of roughly half of these were differentially expressed in the maize mutants (Supplemental Dataset S8). However, many of these were affected even in maize *tha1* and Arabidopsis *psad1*, indicating that these are indirect effects of plastid gene expression defects. In a third study, terms related to the chloroplast nucleoid and chloroplast RNA processing were over-represented among the up-regulated genes in mutants with partial defects in chloroplast or chloroplast and mitochondrial tRNA metabolism (Leister and Kleine, 2016), similar to our observations in maize. Shared up-regulated genes include many PEP-associated proteins, RpoTp/NEP and the plastid tRNA processing enzyme CPZ. Most of these were also up-regulated in maize *tha1* and under high light stress in Arabidopsis (Leister and Kleine, 2016) (Supplemental Dataset S8), indicating that these result from a defect in photosynthesis or protein homeostasis, which are indirect effects of disrupted plastid gene expression.

In summary, the strongest conserved trends in maize and Arabidopsis involve down-regulation of PhANGs in the absence of plastid translation, and up-regulation of genes encoding PEP-associated proteins and several other plastid gene expression factors when photosynthesis is disrupted. The data also suggest conserved up-regulation of cytosolic ribosome biogenesis genes in seedlings lacking plastid ribosomes, which we show is associated with elevated TOR activity in maize (see above). By contrast, there was little indication of conserved responses for genes in retrograde Clusters 2 and 3, and the up-regulation of genes involved in chloroplast translation and RNA metabolism (primarily in retrograde Cluster 7) was weak in the Arabidopsis datasets. These differences may reflect divergent regulatory mechanisms, or they may be due to differences in the tissues analyzed (cotyledon *versus* leaf) and growth conditions employed (e.g. sucrose-supplemented growth medium *versus* growth in soil).

### Extensive retrograde control of transcription factor expression

Our differentially expressed gene set contains at least 113 genes encoding transcription factors (Supplemental Dataset S5). Orthologs of several of these have been highlighted in studies of retrograde signaling in Arabidopsis, including GLK1 paralogs discussed above. Similar to GLK1/2, the paralogous transcription factors GNC and CGA1 are positive regulators of chloroplast development in Arabidopsis (reviewed in Zubo et al., 2018; Cackett et al., 2021). Our differentially expressed gene set includes two putative orthologs of GNC and CGA1. Interestingly, one is up regulated in all four mutants whereas the other is down regulated in the three mutants with defective plastid transcription/translation (Figure 3B, Supplemental Dataset S5). Arabidopsis CGA1 is strongly up-regulated in the *pap7* and *psad1* mutants (Glässer et al., 2014; Grübler et al., 2021) (Supplemental Dataset S5), similar to its Cluster 9 homolog in maize. These observations suggest that the current view of CGA1 function may be overly simplistic.

The paralogous transcription factors CIA2 and CIL activate genes encoding plastid ribosomal proteins and plastid import early in chloroplast development (Sun et al., 2009; Gawronski et al., 2021; Li et al., 2021b; Yang et al., 2022). A maize homolog (Zm00001d014664) was down-regulated in all four mutants, consistent with reduced expression of Arabidopsis CIA2 in norflurazon-treated seedlings (Supplemental Dataset S5). Interestingly, CIA2’s down-regulation is accompanied by up-regulation of the chloroplast biogenesis genes it is proposed to activate (Figure 3). Several myb transcription factors are notable for their very strong down-regulation in some or all of our mutants (Supplemental Dataset S5): one is orthologous to Arabidopsis AT2G38090, which exhibits similar retrograde regulation (Grübler et al., 2021), whereas the others have not, to our knowledge, been connected previously with retrograde signaling or chloroplast development. Other differentially expressed transcription factors provide links between retrograde signaling and many other signaling pathways. Examples include orthologs of PhyA and PIF3/4 (light signaling), orthologs of ARR10, ARF6, ARF16, ETR2 and EIN4 (cytokinin, auxin, ethylene signaling), and three WRKY transcription factors (abiotic stress, sucrose signaling) (Supplemental Dataset S5).

In summary, differentially expressed transcription factors in the maize mutants include some that were previously linked to chloroplast biogenesis and retrograde signaling, and many others that we have not seen discussed in this context. These include transcription factors involved in development, light signaling, energy signaling, and hormone responses, indicating that a complex interplay among regulatory networks underlies the transcriptional reprogramming triggered by disrupted chloroplast biogenesis. The polarity of changes in expression of key transcription factors is often conserved in analogous Arabidopsis treatments, but does not always correlate with that of genes these factors are proposed to regulate, indicating complexities that are not understood.

### The Zm-*ptac12* transcriptome suggests that Zm-PTAC12 does not play a direct role in either phytochrome or retrograde signaling

The Zm-*ptac12* transcriptome provided an opportunity to assess whether the role in phytochrome signaling attributed to Arabidopsis HMR/PTAC12 is conserved in maize. Arabidopsis *hmr* mutants have elongated hypocotyls when grown in red light, a phenotype that motivated experiments showing that HMR promotes phytochrome signaling by triggering the degradation of Phytochrome Interacting Factors (PIFs) (Chen et al., 2010; Hernandez-Verdeja and Strand, 2018). However, we have not observed morphological differences between Zm-*ptac12* and maize mutants lacking other PEP-associated proteins (Williams-Carrier et al., 2014), so we questioned whether the role of PTAC12/HMR in phytochrome signaling is conserved in maize.

A role for Zm-PTAC12 in light-signaling should be detectable as changes in the Zm-*ptac12* transcriptome that are not shared by the other maize mutants. The Zm-*ptac12* and *tha1* transcriptomes were very similar (Pearson correlation ∼0.95) (Figure 2, Supplemental Figure S1), and a direct comparison of the Zm-*ptac12* and *tha1* transcriptomes by DESeq2 produced 323 differentially-expressed genes (Supplemental Dataset S9). Where differences were observed, Zm-*ptac12* usually resembled Zm-*murE* and *ppr5,* suggesting that these result from the considerable loss of plastid ribosomes in the Zm-*ptac12* mutant. We identified only 43 genes in the differentially expressed set whose fold-change in Zm-*ptac12* was substantially different from that in either *tha1* or *ppr5/*Zm-*murE*, and these have no apparent connection to light-regulated networks or chloroplast biology (Supplemental Dataset S9). We also compared our data to the 106 HMR-dependent PIF direct target genes in Arabidopsis (Qiu et al., 2015), which were identified from analysis of plants grown in continuous red light. The ortholog of only one of those genes was differentially-expressed in the *tha1* versus Zm-*ptac12* comparison, whereas orthologs of 24 of those genes were differentially expressed in multiple maize mutants indicating that altered expression results from retrograde signaling in maize (Supplemental Dataset S9). These results strongly suggest that Zm-PTAC12 does not directly affect light signaling in maize seedlings grown in diurnal cycles. HMR has also been suggested to directly mediate retrograde signaling via its dual functions in chloroplast and nuclear transcription (Tadini et al., 2020; Grübler et al., 2021). Again, the strong similarities between the Zm-*ptac12* transcriptome and those of the other mutants we analyzed argue against this possibility in maize.

## DISCUSSION

Almost all of the research into mechanisms underlying plastid-to-nucleus retrograde signaling has been done in Arabidopsis. The emphasis of the biogenic signaling literature has been on the retrograde regulation of PhANGs, and there is strong evidence that light signaling, tetrapyrroles, the transcription factor GLK1, and the chloroplast-localized protein GUN1 are relevant to decreased PhANG expression in seedlings with global defects in plastid transcription and translation (reviewed in Chan et al., 2016; Kleine and Leister, 2016; Larkin, 2016; Hernandez-Verdeja and Strand, 2018; Jiang and Dehesh, 2021; Wu and Bock, 2021). Our examination of retrograde signaling through a different lens highlighted features that have received little attention, and incorporated previously disparate observations into a model that places retrograde signals and the transcriptomic consequences of plastid dysfunctions into a developmental context.

The transcriptome remodeling in the four mutants we analyzed presented in two primary patterns or syndromes. The “chlorotic” syndrome is displayed by the Zm-*ptac12* and *tha1* transcriptomes, whose similarity suggests that Zm-PTAC12 does not directly coordinate nuclear and chloroplast gene expression or mediate phytochrome signaling, as has been proposed for its Arabidopsis ortholog (Chen et al., 2010; Hernandez-Verdeja and Strand, 2018; Tadini et al., 2020; Grübler et al., 2021). The albino syndrome is displayed by the Zm-*murE* and *ppr5* mutants. PhANG repression is a feature of the albino syndrome but not the chlorotic syndrome, consistent with the paradigm developed in Arabidopsis and with prior studies in the grasses (Mayfield and Taylor, 1984; Batschauer et al., 1986; Burgess and Taylor, 1987; Rapp and Mullet, 1991; Hess et al., 1994; Li et al., 2017; Wang et al., 2020). However, PhANGs make up only a minority of the genes with this type of retrograde response, and the others are involved in diverse processes both within and outside plastids. Furthermore, the vast majority of genes encoding proteins involved in chloroplast biogenesis are not coregulated with PhANGs; instead, they are generally up-regulated in the mutants (Figure 3A), along with many genes encoding extra-plastidic proteins (Supplemental Dataset S2, Retrograde Clusters 7-10). In accord with these wide-ranging effects, the expression of more than 100 transcription factors was altered in these mutants, including many that had not previously been linked to retrograde signaling or chloroplast development (Supplemental Table S5).

Genes whose expression increases in response to disrupted chloroplast biogenesis have been reported previously (e.g. Biehl et al., 2005; Woodson et al., 2013; Kleine and Leister, 2016; Grübler et al., 2017; Wang et al., 2020; Grübler et al., 2021), but aside from NEP, whose expression is well known to increase when PEP is compromised (Tadini et al., 2020), they have received little attention with regard to their functions or mechanisms of up-regulation. Roughly half of the genes in our differentially-expressed set were expressed at higher levels in mutants than in the wild-type. Our data suggest that ectopically-active TOR accounts for enhanced expression of cytosolic ribosome biogenesis and cell proliferation genes in mutants lacking plastid ribosomes, whereas disrupted photosynthesis leads to increased expression of many genes involved in early stages of chloroplast biogenesis. The correspondence of gene sets that cluster based on their expression patterns in the four mutants (retrograde regulons) and those that cluster based on expression dynamics along the developmental gradient of the leaf blade (developmental regulons) demonstrates a clear link between retrograde signals and gene regulation during chloroplast development. The results provide strong support for the integration of multiple retrograde signals into the normal program of photosynthetic differentiation. These key themes that emerged from our analyses are discussed below.

### Retrograde signals as central actors during normal chloroplast development

The developmental dynamics of genes in each retrograde cluster were remarkably uniform (with the exception of Cluster 3), whereas genes in different retrograde clusters were characterized by distinct developmental dynamics (Figure 5). Almost all of the up-regulated genes peak in expression prior to the sink-source transition, whereas those that were down-regulated peak after the sink-source transition. Both the up- and down-regulated gene sets fall into several classes based on the magnitude of the response in particular mutants, and these different classes map onto distinct developmental regulons with subtle differences in the steepness of the expression gradient or the location of peak expression (Figure 5). For example, among the up-regulated retrograde clusters, the expression of genes in Cluster 8 peaks earliest in development, followed by those in Cluster 9/10 and then Cluster 7. This concordance between various retrograde responses and distinct developmental dynamics strongly suggests that the same retrograde signals that are impacted by chloroplast dysfunctions in the mutants are integral components of the regulatory circuits underlying normal photosynthetic differentiation.

In a study of the virescent *cue8* mutant in Arabidopsis, the similarity of several molecular and cell biological phenotypes to those at early stages of chloroplast development led to the suggestion that decreased PhANG expression in plants with defective plastids was not due to their repression, but rather to failure to reach the developmental stage during which PhANG activation occurs (Loudya et al., 2020). Our data provide strong support for this view. More than one thousand genes that are expressed early in normal chloroplast biogenesis were expressed at elevated levels in the mutants, whereas a similar number of genes that are expressed late in normal chloroplast biogenesis were down-regulated in the mutants. Furthermore, comparison of each mutant transcriptome to a developmental series along a wild-type leaf blade provided evidence that the absence of plastid ribosomes/PEP-mediated transcription halts development at a stage that slightly precedes the sink-source transition, whereas the failure to achieve photosynthetic competence stalls the developmental program at a slightly later stage. These results suggest that the sweeping changes in gene expression observed when chloroplast biogenesis is inhibited are not compensatory, as is sometimes suggested, but rather are an emergent property of disrupting retrograde signals that play key roles during normal chloroplast development.

### Positive and negative retrograde signals as coordinators of photosynthetic differentiation

Analyses of “genomes uncoupled” (*gun*) mutants (Susek et al., 1993) led to the proposal that Mg-protoporphyrin IX is a negative signal leading to PhANG repression (reviewed in Strand et al., 2003). However, more recent data provide compelling evidence for an alternative view - that a heme-related molecule activates PhANGs (Woodson et al., 2011; Terry and Smith, 2013; Woodson et al., 2013; Larkin, 2016; Wu and Bock, 2021). An attractive feature of the heme model is that it provides a clear link between plastid gene expression and PhANG expression. Plastid-encoded tRNA^glu^ is the precursor for tetrapyrroles synthesized in the plastid (Schön et al., 1986) and is transcribed by PEP (Kanamaru et al., 2001; Williams-Carrier et al., 2014), which is synthesized by plastid ribosomes. Therefore, loss of plastid ribosomes results in loss of tRNA^glu^ and a decrease in heme synthesis. Indeed, it has been shown that PhANG expression is closely tied to PEP activity (Woodson et al., 2013; Diaz et al., 2018).

In a converging line of evidence, a positive biogenic signal was recently inferred based on transcriptome dynamics in two systems: light-induced greening in an Arabidopsis single-cell culture (Dubreuil et al., 2018) and the developmental gradient of the wheat leaf (Loudya et al., 2021). In both contexts, PhANG activation follows shortly after activation of PEP-mediated transcription, suggesting that PEP activity produces a positive signal that activates PhANGs. Likewise, during maize leaf development, expression of genes encoding proteins that are required for PEP activity and the up-regulation of plastid genes that are transcribed by PEP precedes the up-regulation of PhANGs (Figure 3) (Wang et al., 2014; Chotewutmontri and Barkan, 2016). These observations support the model that a heme-derived molecule is a positive signal that activates PhANGs during chloroplast development (Terry and Smith, 2013; Shimizu et al., 2019) in both monocots and dicots.

The transcription factor GLK1 has been proposed to activate PhANGs in response to a biogenic retrograde signal (Leister and Kleine, 2016; Martin et al., 2016; Hernandez-Verdeja and Strand, 2018; Grübler et al., 2021). Our results indicate, however, that the quantitative relationships between PEP activity, PhANG expression and GLK1 expression are non-linear. The expression of most PhANGs was minimally affected in the Zm-*ptac12* mutant despite its large deficit in plastid rRNA (Figure 1) and tRNA^glu^ (Williams-Carrier et al., 2014). Furthermore, whereas GLK1 expression was reduced in the albino mutants, it was also reduced in non-photosynthetic mutants with normal plastid gene expression (maize *tha1,* Arabidopsis *psad1*), unlike the PhANGs it regulates. These features suggest that threshold concentrations and/or cooperative interactions among signaling components are involved in retrograde regulation of PhANGs.

Our data also provide evidence for a second plastid-derived signal that feeds into the chloroplast biogenesis program - a negative biogenic signal that acts later in chloroplast development to shut down chloroplast biogenesis genes when chloroplasts approach maturity. Our differentially-expressed gene set included at least 170 genes involved in chloroplast biogenesis (plastid gene expression, division, protein targeting, assembly of photosynthetic complexes), almost all of which were expressed at elevated levels in the mutants (Figure 3A, Supplemental Dataset S5). Expression of these genes peaks in a “chloroplast biogenesis zone” prior to the sink-source transition during normal leaf development, and drops to low levels thereafter (Figure 3A). The correlated developmental timing of the sink-source transition and repression of chloroplast biogenesis genes, in conjunction with elevated expression of these genes in our mutants, provides evidence for a repressive signal that is produced in response to some aspect of chloroplast function.

A clue about the nature of the repressive signal(s) comes from the fact that up-regulation of many chloroplast biogenesis genes was observed across all mutants in our study. These include many genes involved in early steps in chloroplast biogenesis, such as PEP-associated proteins, NEP, proteins involved in plastid tRNA/rRNA processing, plastid protein import and plastid division (e.g. ARC5, FtsZ1) (Figure 3, Supplemental Dataset 5). Up-regulation of genes encoding NEP and PEP-associated proteins is also evident in Arabidopsis norflurazon, lincomycin, *psad1, mterf6* and *prors1* transcriptomes (Supplementary Dataset S5). Furthermore, proteome analysis of chlorotic Clp protease mutants in Arabidopsis (Nishimura and van Wijk, 2015) revealed increased abundance of many proteins involved in chloroplast biogenesis (Nishimura and van Wijk, 2015), mirroring the transcriptome data for our maize mutants.

A parsimonious explanation of these observations is that photosynthesis produces the repressive signal for the chloroplast biogenesis gene set, via, for example, exported sugars or reducing equivalents. This could potentially involve energy signaling, given that the expression of numerous SnRK1 regulators is altered in all four mutants (e.g. KING1 and many trehalose-6-phosphate synthase and FLZ/DUF581 genes) (Supplementary Dataset S2). An alternative possibility is that the disruption of chloroplast protein homeostasis triggers increased expression of biogenesis genes. However, we favor the hypothesis that a negative signal derived from photosynthesis is the primary driver of the retro-regulation of the biogenesis gene set for two reasons: (i) the repression of chloroplast biogenesis genes coincides with the sink-source transition during normal leaf development, and (ii) homologs of the heat shock transcription factor proposed to underlie the chloroplast unfolded protein response in Arabidopsis (Llamas et al., 2017) are not represented among the up-regulated genes in our data. Several operational signals (singlet oxygen, sugars, thylakoid redox) have been proposed to feed into what is typically referred to as “biogenic” signaling (Tadini et al., 2012; Glässer et al., 2014; Page et al., 2017). Our findings further blur the line between operational and biogenic signaling by providing evidence that photosynthesis itself represses the chloroplast biogenesis program during photosynthetic differentiation.

### A connection between the buildup of the chloroplast genetic system, TOR, and exit from the cell proliferation phase of leaf development

TOR regulates cell proliferation and metabolism in response to nutrient and energy status (Brunkard, 2020), so effects of photosynthetic defects on TOR activity are to be expected. However, we had not anticipated the nature of the TOR-chloroplast crosstalk revealed by our data. Genes involved in cytosolic ribosome biogenesis, translation, and cell proliferation, which are activated by TOR (Xiong et al., 2013; Scarpin et al., 2020), are prominent among the gene set that is most strongly up-regulated in the two albino mutants (retrograde Cluster 8 in Figure 2, Figure 3B). Expression of these genes in the wild-type peaks in the cell proliferation zone at the leaf base (Figure 3B), correlating, as expected, with a marker of TOR activity (eS6-S240P) (Figure 4A). However, contrary to our naïve expectation, both of these readouts of TOR activity indicated elevated TOR activity in the mutants (Figure 3B, Figure 4B). Furthermore, these TOR activity markers correlate with the magnitude of the plastid ribosome deficiency: highest in the two albino mutants, somewhat lower in Zm-*ptac12* and minimal in the *tha1* mutant.

We propose a simple model to account for these correlations that derives from the fact that TOR is positively regulated by free amino acids and nucleotides (Brunkard, 2020): the massive diversion of amino acids and nucleotides toward the developing chloroplast following PEP activation transiently represses TOR activity. Thus, mutants that cannot activate PEP (such as Zm-*murE)* or make plastid ribosomes (such as *ppr5)* maintain high TOR activity beyond the normal cell proliferation zone. According to this hypothesis, the “retrograde signal” underlying the altered expression of ribosome biogenesis/cell proliferation genes does not emanate from plastids. Instead, it is the failure of the defective plastids to consume nutritional resources that accounts for this effect.

These observations raise the possibility that the cessation of cell proliferation early in leaf development is causally connected with the up-regulation of PEP-mediated transcription in chloroplasts. Related to this point, norflurazon prevents the transition from cell proliferation to cell expansion and increases cell number in expanding leaves in Arabidopsis (Andriankaja et al., 2012; Van Dingenen et al., 2016). On this basis, it was proposed that photosynthetically-active chloroplasts emit retrograde signals that promote the transition from cell proliferation to cell expansion. Our data align nicely with those observations but reframe their interpretation.

Norflurazon prevents plastid ribosome buildup, mimicking the dysfunction of our albino maize mutants. Similarly, Arabidopsis *clb5* mutants, which lack plastid transcripts synthesized by NEP (and therefore lack plastid ribosomes) show an expansion of the zone expressing the cell-proliferation marker CYCB1 (Avendaño-Vazquez et al., 2014). These concordant observations in maize and Arabidopsis suggest that consumption of amino acids and nucleotides for the build-up of the chloroplast translation machinery leads to a transient decrease in TOR activity, thereby promoting exit from the cell division phase of leaf development in both monocots and dicots. We expect that TOR activity would increase once again in mature non-dividing leaf cells, where photosynthesis more than compensates for resources consumed by the chloroplast and is positively correlated with TOR activity (Mohammed et al., 2018; Brunkard et al., 2020).

The complementary developmental dynamics of two phosphorylated eS6 isoforms, eS6-237P and eS6-240P (Figure 4) is intriguing in light of these proposed transitions in TOR activity. eS6-240P abundance declines from base to tip, in concert with an increase in eS6-237P. Furthermore, eS6-237P was not affected by treatment with AZD-8055 (Supplemental Figure S4), a TOR inhibitor that inhibits eS6-240P (Dobrenel et al., 2016). eS6 can be phosphorylated at many additional sites (Williams et al., 2003), and the bulk eS6-237P isoform migrates more rapidly in SDS-PAGE than does the major eS6-240P isoform (Figure 4, and Williams et al., 2003), suggesting that this isoform includes multiple phosphorylated residues. We speculate that this transition between different eS6 phospho-isoforms during photosynthetic differentiation results from transition between TOR signaling modes that occurs in response to the transition of the plastid compartment from sink to source.

### Integration of three plastid-related signals into the program of photosynthetic differentiation: a working model

Our results, when interpretated in the context of the transcriptomic, physiological, and cell biological changes that accompany the proplastid-to-chloroplast transition (Wang et al., 2014; Loudya et al., 2021), lead to a working model involving roles for three plastid-related signals during photosynthetic differentiation (Figure 7). (i) A positive signal derived from heme activates PhANG transcription after the onset of PEP-mediated transcription, as proposed previously (Woodson et al., 2011; Terry and Smith, 2013; Woodson et al., 2013; Dubreuil et al., 2018). This signal would coordinate the synthesis of plastid- and nucleus-encoded subunits of the photosynthetic apparatus, and would trigger changes in metabolism and morphology that collaborate with photosynthesis. (ii) A negative signal derived from photosynthesis represses chloroplast biogenesis genes shortly after the sink-source transition. This would comprise a negative feedback loop, shutting down the chloroplast biogenesis program once it is no longer needed. (iii) The depletion of amino acids and nucleotides during the build-up of the chloroplast gene expression system represses TOR activity, and thereby promotes exit from the cell proliferation phase of leaf development.

**Figure 7.**
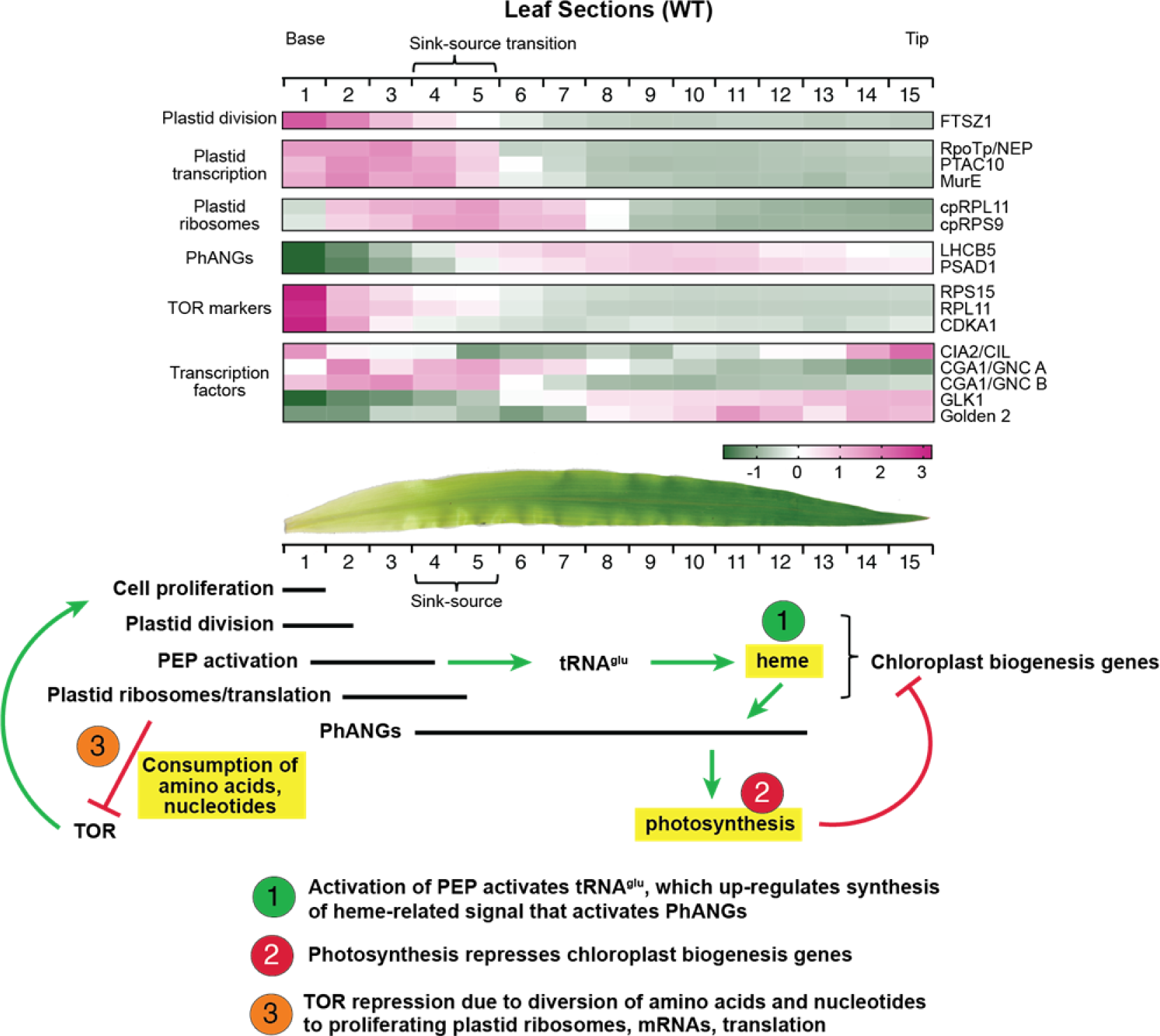
Working model for the integration of three plastid-related signals into the photosynthetic differentiation program. The expression dynamics of marker genes for different phases of photosynthetic differentiation are shown at top (data taken from (Wang et al., 2014)). The peak activity of genes representing the major phases are diagrammed below, together with the proposed integration of three plastid-related signals into this regulatory cascade. Activation of PEP up-regulates tRNA^glu^/heme which activates PhANGs. Genes in the heat map have the following gene identifiers (B73 RefGen v4 nomenclature): FTSZ1-Zm00001d001939, RpoTp/NEP-Zm00001d046835, PTAC10-Zm00001d012141, MurE-Zm00001d047128, cpRPL11-Zm00001d027421, cpRPS9-Zm00001d034192, LHCB5-Zm00001d007267, PSAD1-Zm00001d034543, RPS15-Zm00001d018979, RPL11-Zm00001d003896, CDKA1-Zm00001d027373, CIA2/CIL-Zm00001d014664, CGA1/GNC-Zm00001d046354 and Zm00001d016361, GLK1-Zm00001d044785, Golden2-Zm00001d039260.

The behavior of retrograde Cluster 7 suggests that some genes respond to multiple retrograde signals. Cluster 7 is strongly enriched for functions in chloroplast biogenesis - in particular for genes involved in chloroplast translation and RNA metabolism. These genes are strongly up-regulated in the *tha1,* Zm- *ptac12* pair, consistent with a repressive signal arising from photosynthesis. That said, they are less strongly up-regulated in the two albino mutants, indicating that a photosynthesis-related signal is insufficient to account for their behavior. It is possible, however, that the intersection of a heme-related activating signal and a photosynthesis-related repressive signal underlies the behavior of genes in Cluster 7. According to this view, the albino mutants lack the activating signal and thus fail to up-regulate this gene set. An alternative view of the same phenomenon is that chloroplast development in the albino mutants is stalled prior to the stage at which expression of the plastid translation/RNA metabolism gene cohort peaks. The responses of genes in retrograde Clusters 1, 2, and 6 (down regulated in all four mutants, but to varying degrees) might likewise result from an integration of the positive and negative signal on these gene sets.

Three paralogous pairs of transcription factors have been closely tied to chloroplast development: G2/GLK1, CIA2/CIL, and CGA/GNC (reviewed in Cackett et al., 2021). The expression dynamics of these genes, when compared with those of marker genes for successive stages of chloroplast development (Figure 7), offers clues about their place in the regulatory cascade. Among these, the early expression of CIA2/CIL makes it a good candidate for initiating the chloroplast development program at the leaf base. This possibility is supported by the decreased expression of genes involved in plastid import and chloroplast translation in *cia2/cil* mutants in Arabidopsis (Sun et al., 2009; Li et al., 2021b; Yang et al., 2022). The expression dynamics of maize orthologs of CGA and GNC (Figure 7) are consistent with the possibility that they are up-regulated by CIA2/CIL, and that they, in turn, up-regulate genes involved in PEP-mediated transcription and plastid translation. The expression dynamics of G2 and GLK1 are similar to that of PhANGs, supporting the current view that they activate PhANGs relatively late in the chloroplast biogenesis program (Waters et al., 2009; Cackett et al., 2021). Each gene in this hypothetical regulatory cascade is impacted by retrograde signals: *g2, glk1, cia2/cil,* and one *cga1/gnc* homolog are down-regulated in our mutants, whereas the other *cga1/gnc* homolog is up-regulated (Figure 3B, Supplementary Dataset S5). Crosstalk between light signaling, retrograde signaling, and chloroplast development features prominently in the Arabidopsis literature. However, early steps in chloroplast development in the grasses proceed in the absence of light (Baumgartner et al., 1989, 1993), so light-retrograde crosstalk, if it exists in the grasses, would come into play toward the end of the cascade.

This working model provides a new framework for the interpretation of existing data, and for the design of experiments to test and elaborate on this general scheme. For example, a major gap in the model concerns the nature of the heme-related and photosynthesis-derived signals.

Clues may come from identifying heme- and photosynthesis-related metabolites whose abundance correlates with transcriptome signatures, both along the length of the seedling leaf blade (Wang et al., 2014) and in chloroplast biogenesis mutants such as those analyzed here.

## MATERIALS AND METHODS

### Plant Material

The mutants analyzed here were described previously: *tha1-1* (Voelker and Barkan, 1995; Voelker et al., 1997), *ppr5-1* (Beick et al., 2008), Zm-*ptac12-2* (Williams-Carrier et al., 2014), Zm-*murE-2* (Williams-Carrier et al., 2014), Zm-*hcf136-1* (Chotewutmontri et al., 2020), *psa3-1* (Shen et al., 2017), *pet2-1/-3* (Williams-Carrier et al., 2010). They arose in a mixed genetic background that included the inbred line B73 and varying contributions from other genotypes. To increase genetic uniformity, the *ppr5-1* mutation was introgressed into B73 four times, Zm-*murE-2* was introgressed into B73 three times, and Zm-*ptac12-2* and *tha1-1* were introgressed into B73 two times. Plants used for RNA-seq analyses of *tha1,* Zm-*ptac12-2,* Zm- *murE,* and *ppr5-*1 were grown in soil in diurnal cycles (16 h days at 100 µE/m^2^/s, 28°C) and 8 h nights at 25°C for 8-9 days, until the third leaf just started to emerge. The second leaf to emerge was harvested 2 h into the light cycle by cutting at the point of blade expansion above the whorl, close to the ligule of leaf 1. Replicates used total leaf RNA from three different individual mutant seedlings. Three phenotypically normal siblings were pooled from each mutant line, and each of these pools served as one “wild-type” replicate. A fifth wild-type replicate used RNA from a phenotypically-normal sibling of a different mutant that had been grown in parallel with the first replicate samples. RNA-seq data from mutants lacking either PSII, PSI, or cytochrome *b_6_f* (*hcf136-1, psa3-1, pet2-1/-3,* respectively) had been generated for a different project, but were used here as a point of comparison. These seedlings were grown in soil in cycles of 16 h of light (300 µE/m^2^/s) at 28 °C and 8 h dark at 26 °C for 8 days until leaf 3 started to emerge. On day 8, plants were shifted to the dark for 1 hour at midday, and then reilluminated for 15 min. Leaves 2 and 3 were then harvested by cutting at the point of leaf expansion, just above the ligule of leaf 1. Two biological replicates involving different seedlings from the same planting were generated.

Immunoblot analyses of eS6 phosphorylation along the leaf gradient was performed with material prepared as described in the study of transcriptome dynamics we used as the point of comparison (Wang et al., 2014). The inbred line B73 was grown in diurnal cycles: 12 h light (250 µE/m^2^/s at 31 °C) and 12h dark at 22 °C. The third leaf to emerge was harvested on the 13th day after planting, and was divided into 15 segments of equal length. Segments from 3 seedlings were pooled for each sample. Immunoblot analysis of eS6 phosphorylation in the mutants used tissue prepared and harvested in the same manner as that used for RNA-seq (see above).

Plants were treated with the TOR inhibitor AZD-8055 as described in (Salazar-Diaz et al., 2021) with minor modifications. Maize seedlings were germinated in moist paper towels and watered with 50 µM AZD-8055 (or mock treated) at midday on days 5 and 7. The indicated section of leaf 2 was harvested at midday on day 8. Each sample was pooled from three seedlings. Arabidopsis Col-0 was grown in long day conditions (16h day at 100 µE/m^2^/s, 8 h dark) on MS medium supplemented with 1%sucrose. Four hours after dawn on day 14, each seedling was transferred to a well of a 24-well plate filled with 500 µl of MS medium supplemented with 1% sucrose and 2 µM AZD-8055. The plate was placed in light (100 µE/m^2^/s) for 4 h, then 1 h dark before the rosette leaves were harvested for dark sample. The re-illuminated samples were harvested after re-illumination of the plants for 35 min. Each sample was a pool of three plants.

### Immunoblotting

Samples were resolved by SDS-polyacrylamide electrophoresis, electrophoretically transferred to nitrocellulose, and probed with antibodies specific for the proteins indicated in each figure. The antibodies to AtpB, PsbD, and PsaD were raised in rabbits against recombinant proteins. The antibody to RbcL was a gift of William Taylor (formerly of UC-Berkeley). The PetD antibody was purchased from Sigma Genosys. Antibodies to cytosolic ribosomal protein eS6 and its phosphorylated isoforms eS6-S240P and eS6-237P (Enganti et al., 2017) were a generous gift of Albrecht von Arnim (University of Tennessee).

### RNA sequencing and data analysis

Total RNA was treated with Turbo DNase (Ambion, CA) following manufacturer’s recommendations, and polyadenylated RNA was then selected with Sera-Mag Oligo(dT)-Coated Magnetic Particles (Fisher Scientific). Strand-specific RNA-seq libraries were prepared as described by Wang et al. (2011). RNA-seq libraries for *hcf136*, *pet2*, and *psa3* were prepared using the Universal Plus mRNA-Seq library preparation kit with NuQuant (Tecan). Quality trimming at the 3’ end was applied to the first 10 nucleotides of each read and the 5’ adaptor and poly-A tail were removed using CutAdapt version 1.16 (Martin, 2011). Reads were then aligned to the *Zea mays* reference genome (B73 RefGen_v4 assembly) allowing for a maximum of 5% mismatched bases, using STAR version 2.5.3a (Dobin et al., 2013). Raw read counts for each gene that mapped to mRNA regions were obtained using featureCounts (Liao et al., 2014) and the AGPv4.38 genome annotation. Uniquely mapped reads were used for analysis of differential expression by DESeq2 (Love et al., 2014). Each mutant was compared to the same set of five replicate wild-type samples (see above). Genes with a log2-fold change of at least 1.5 and *p*-adj < 0.001 in any of the mutants, as well as an average raw read count of 150 in either the mutant or in WT samples, were defined as differentially expressed.

The gap statistic of the dataset was computed using the clusGap function in R. Clustering was performed using an R implementation of the *k*-means algorithm. We used MapMan (Schwacke et al., 2019) to detect functional categories that were enriched in each gene cluster. RPKM values were computed for all mapped genes in each WT sample. Genes with an average of at least 1 Read per kilobase per million reads mapped (RPKM) across the wild-type samples were assigned to Level 1, 2, and 3 MapMan terms and used to generate a background distribution model for each cluster as described previously (Zones et al., 2015). This background distribution was used to calculate *z*-scores for each MapMan term in each cluster. A MapMan term was defined as significantly overrepresented if its false-discovery rate was < 0.05. Intracellular locations were taken from winner-takes-all annotations for maize from cropPAL (Hooper et al., 2016). Enrichment analysis was performed by permutation as for the MapMan analysis, using genes represented by > 1 RPKM in the WT samples as the background set.

Developmental clusters were calculated based on the maize leaf gradient transcriptomes reported in Wang et al. (2014). The data were scaled such that the distribution of RPKM values at each location along the leaf had 0 mean and unit variance (Supplemental Dataset S6). Next, the developmental dataset was clustered using the same procedure as for our retrograde dataset, with the columns of the matrix used for clustering now representing leaf section rather than genotype. Enrichment analysis was then used to identify developmental clusters that were overrepresented in each of retrograde cluster. For this purpose, we randomly assigned genes with an average of at least 1 RPKM across wild-type samples to a developmental cluster to generate a background distribution model, which gives the number of genes from a given developmental cluster that would be expected to be represented in a given retrograde cluster by chance. This background distribution was used to calculate *z*-scores, which compared the actual number of developmental genes represented in a retrograde cluster with the number expected by chance from the background distribution. A developmental cluster was defined as significantly overrepresented in a given retrograde cluster if its false-discovery rate (FDR) was < 0.05.

### Accession Numbers

Sequence data from this article can be found in the SRA database under BioProject number PRJNA825557.

## Acknowledgements

We are grateful to Tom Brutnell (formerly of the Donald Danforth Plant Sciences Center, St. Louis, MO) for advice on experimental design at the start of this project, Dustin Mayfield-Jones (formerly of the Donald Danforth Plant Sciences Center) for preparing and providing information about the RNA-seq libraries, Rosalind Williams-Carrier (University of Oregon) for technical assistance and comments on the manuscript, Albrecht von Arnim (University of Tennessee) for providing antibodies to eS6 and for helpful discussions, and Meng Chen (University of California) for helpful discussions. This work was supported by grants IOS-1339130 and MCB-2034758 to AB from the US National Science Foundation.

## Author Contributions

RK analyzed data and edited the paper, PC analyzed data, performed research, designed research, and edited the paper, SB performed, designed research, and edited the paper, AB designed research, analyzed data, and wrote the paper.

## Supplemental Data Files

**Supplemental Dataset S1**. Sequencing data summary, including number of mapped reads per library, read counts per gene and complete DESeq2 output.

**Supplemental Dataset S2**. Differentially expressed gene set annotated with fold-change, retrograde and developmental cluster, functional annotations, intracellular locations, Arabidopsis orthologs, and data from corresponding genes in other maize mutants and from various published Arabidopsis datasets (Woodson et al., 2013; Glässer et al., 2014; Grübler et al., 2017; Zhao et al., 2019).

**Supplemental Dataset S3**. Enrichment of cropPal intracellular location terms in Retrograde Clusters

**Supplemental Dataset S4**. Enrichment of MapMan terms in Retrograde Clusters **Supplemental Dataset S5**. Retrograde and developmental data for extracted gene sets parsed by functional categories.

**Supplemental Dataset S6.** Developmental clusters calculated from the leaf gradient transcriptome data of Wang et al (2014).

**Supplemental Dataset S7:** Spearman correlation values for mutant transcriptomes and leaf section transcriptomes.

**Supplemental Dataset S8**. Arabidopsis Core Retrograde Modules (Glässer et al., 2014; Leister and Kleine, 2016; Grübler et al., 2021) annotated to indicate retrograde regulation of maize orthologs.

**Supplemental Dataset S9.** Comparison of Zm-*ptac12* transcriptome to *tha1* and to HMR-dependent PIF targets in Arabidopsis.

---

**Supplemental Figure S1.**
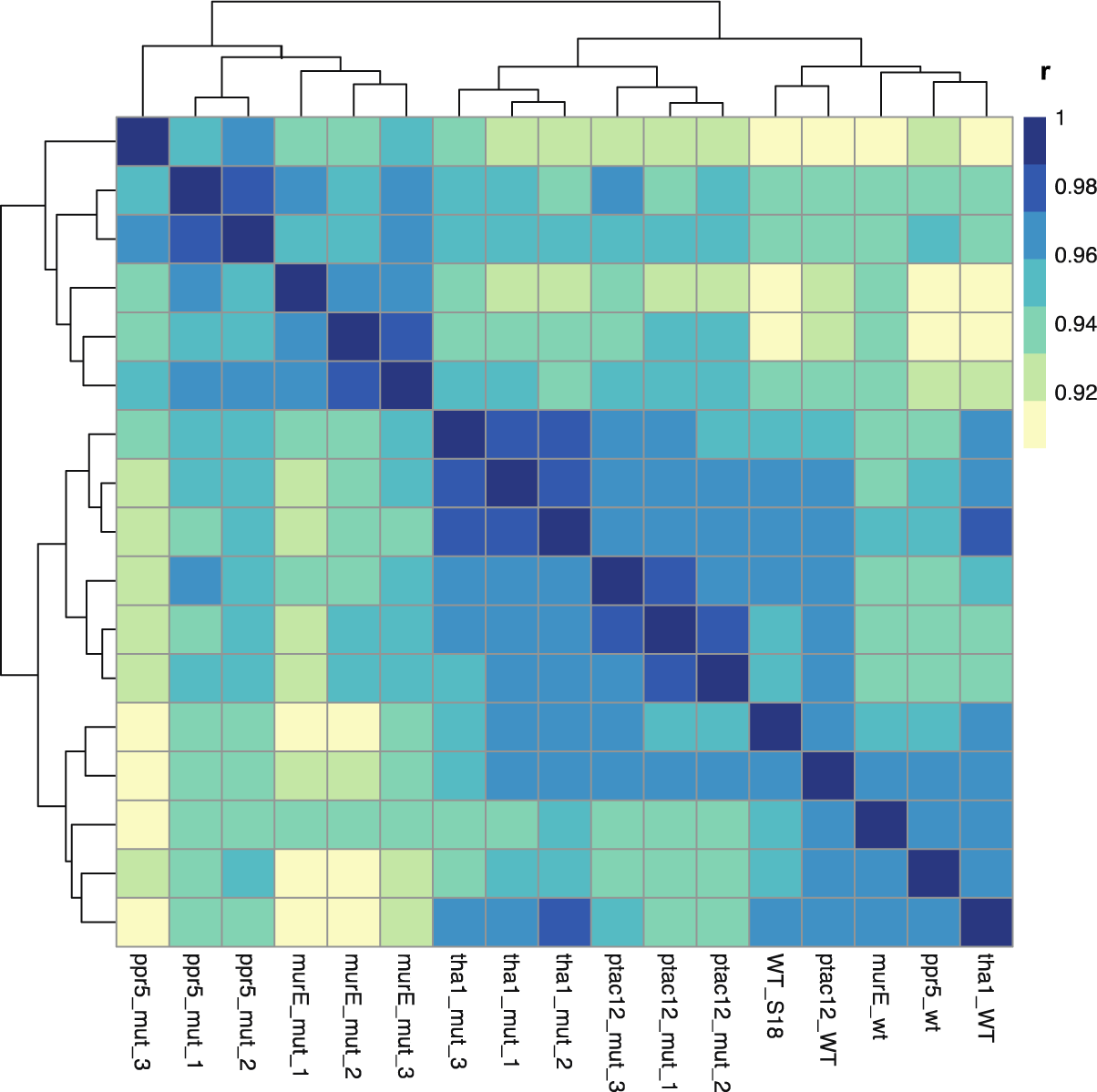
Correlation matrix of RNA-seq datasets. RPKM values were standardized via *z*-scoring and Pearson correlations were computed for pairwise combinations. The Pearson’s distance for each pairwise combination was computed as 1-|Pearson coefficients|. Wild-type (WT) siblings of each mutant comprised four replicate wild-type samples. A fifth wild-type sample (WT_S18) used RNA from a phenotypically-normal sibling of a different mutant that had been grown in parallel with the first replicate samples.

**Supplemental Figure S2.**
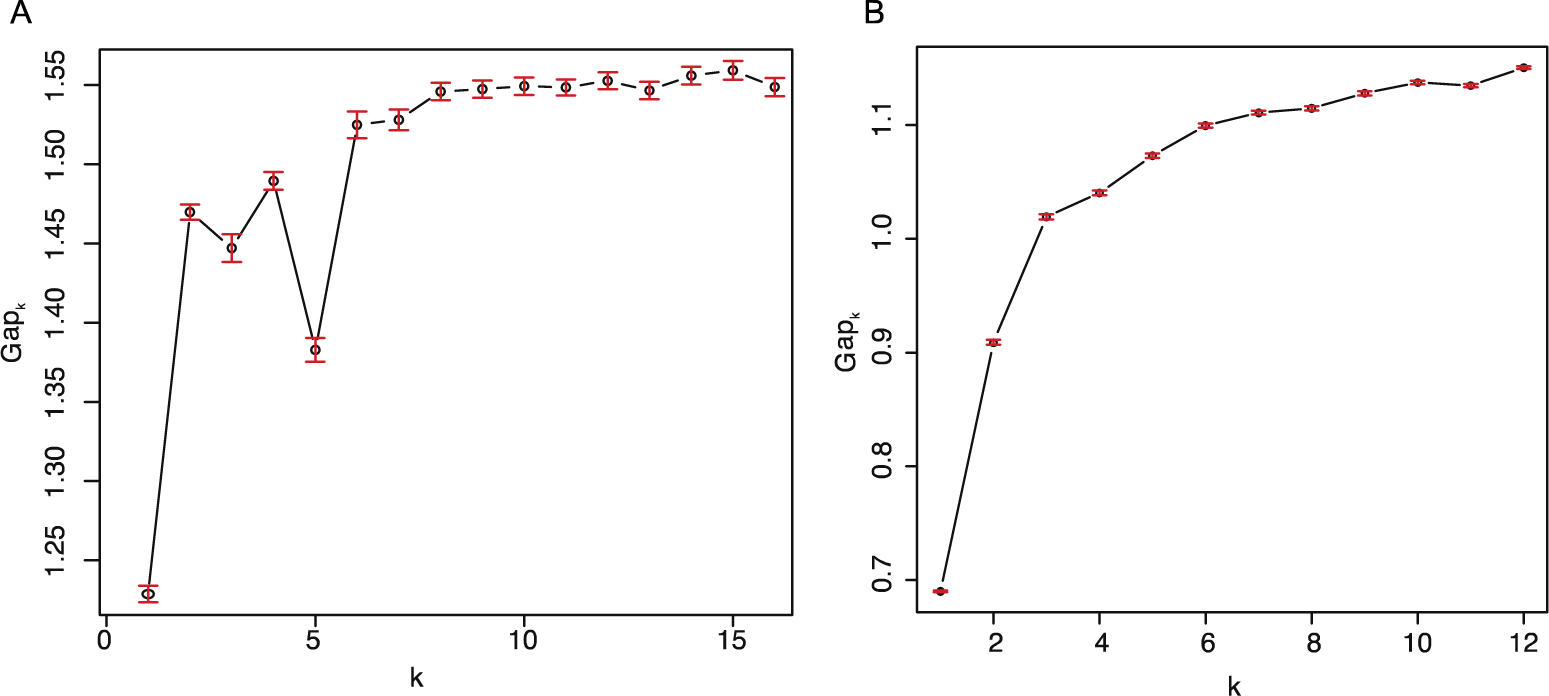
Gap statistic analysis for *k*-means clustering. The gap statistic (Tibshirani et al., 2001) was calculated using the clusGap function in R to determine the optimal number of clusters for (A) differentially expressed genes reported here and (B) transcriptomes of fifteen leaf sections along the maize seedling leaf blade reported by Wang et al. (2014).

**Supplemental Figure S3.**
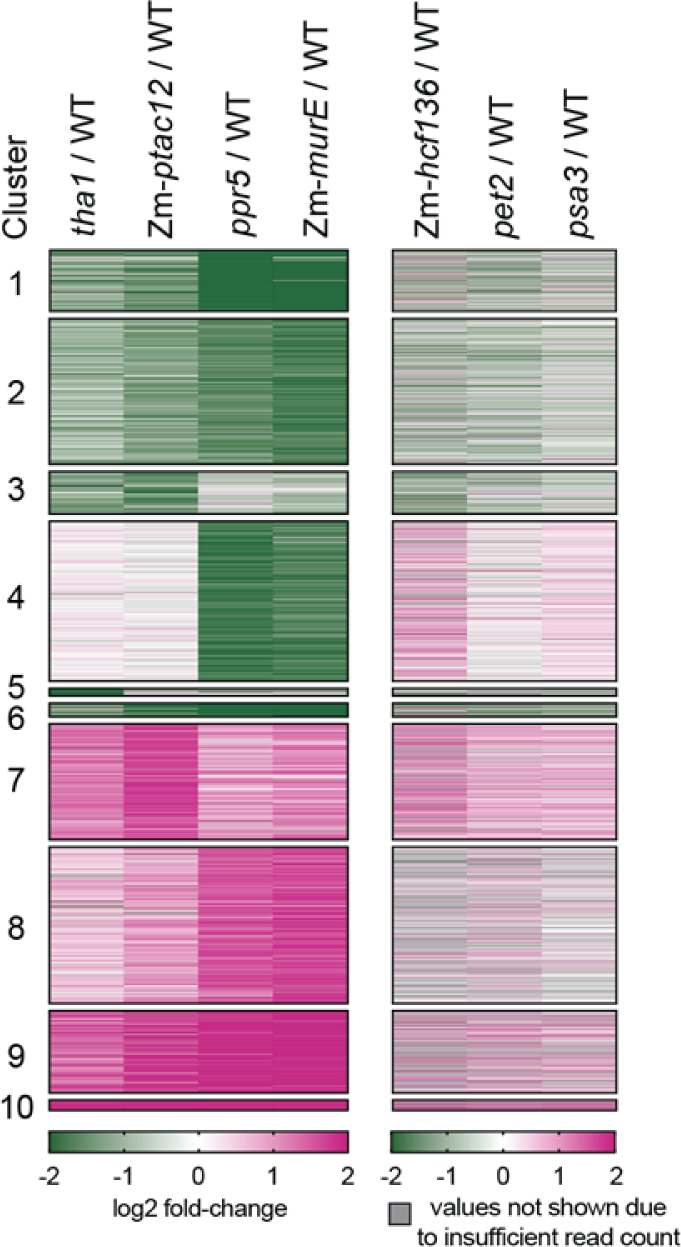
Effects of specific photosynthetic defects on expression of genes in the differentially-expressed set. Left: Heat map of differentially expressed genes in this study, as in Figure 2A. Right: Heat map of RNA-seq data from maize mutants lacking either PSII (Zm-*hcf136),* the cytochrome *b_6_f* complex (*pet2*), or PSI (*psa3*) that were collected in the context of a different study. These mutants were grown under different conditions and the samples were sequenced to lower depth than those of the primary mutants analyzed here (see Materials and Methods), so these data are presented only to illustrate trends. Data in the heat map to the right are displayed for genes with an average of at least 50 reads in either the wild-type (WT) or mutant sample.

**Supplemental Figure S4.**
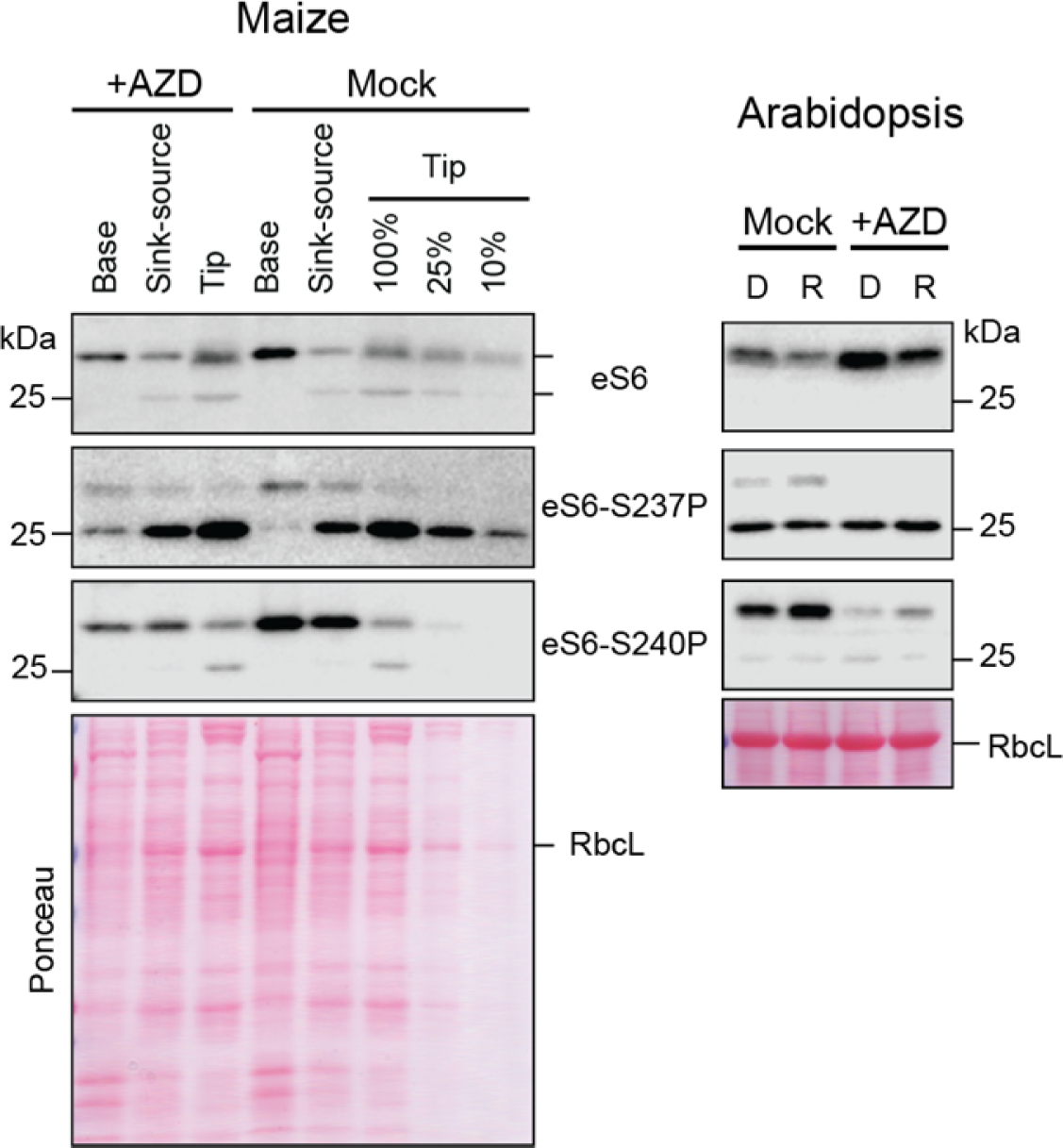
Effects of the TOR inhibitor AZD-8055 on abundance of eS6-S237P and eS6-S240P in maize and Arabidopsis. Maize seedlings (inbred line B73) were treated with AZD-8055 for three days prior to harvest at midday on day 8. Three segments of Leaf 2 were harvested: Base (soil level to 2 cm above), Sink-source (from 2-cm above the soil to the position of ligule of leaf 1), and Tip (apical 2 cm). Samples from three seedlings were pooled. Arabidopsis seedlings (Col-0) were treated with AZD-8055 for 4 hours in the light, then shifted to the dark for 1 h (D) and reilluminated for 35 min (R). Blots were probed with antibodies to eS6 or its phosphorylated isoforms S237P and S240P. Ponceau S-stained filters are shown below to illustrate relative sample loading. The bulk of the eS6-S237P isoform migrates at ∼25 kDa whereas the bulk of the eS6-S240P isoform migrates at ∼28 kDa in both maize and Arabidopsis.

**Supplemental Figure S5.**
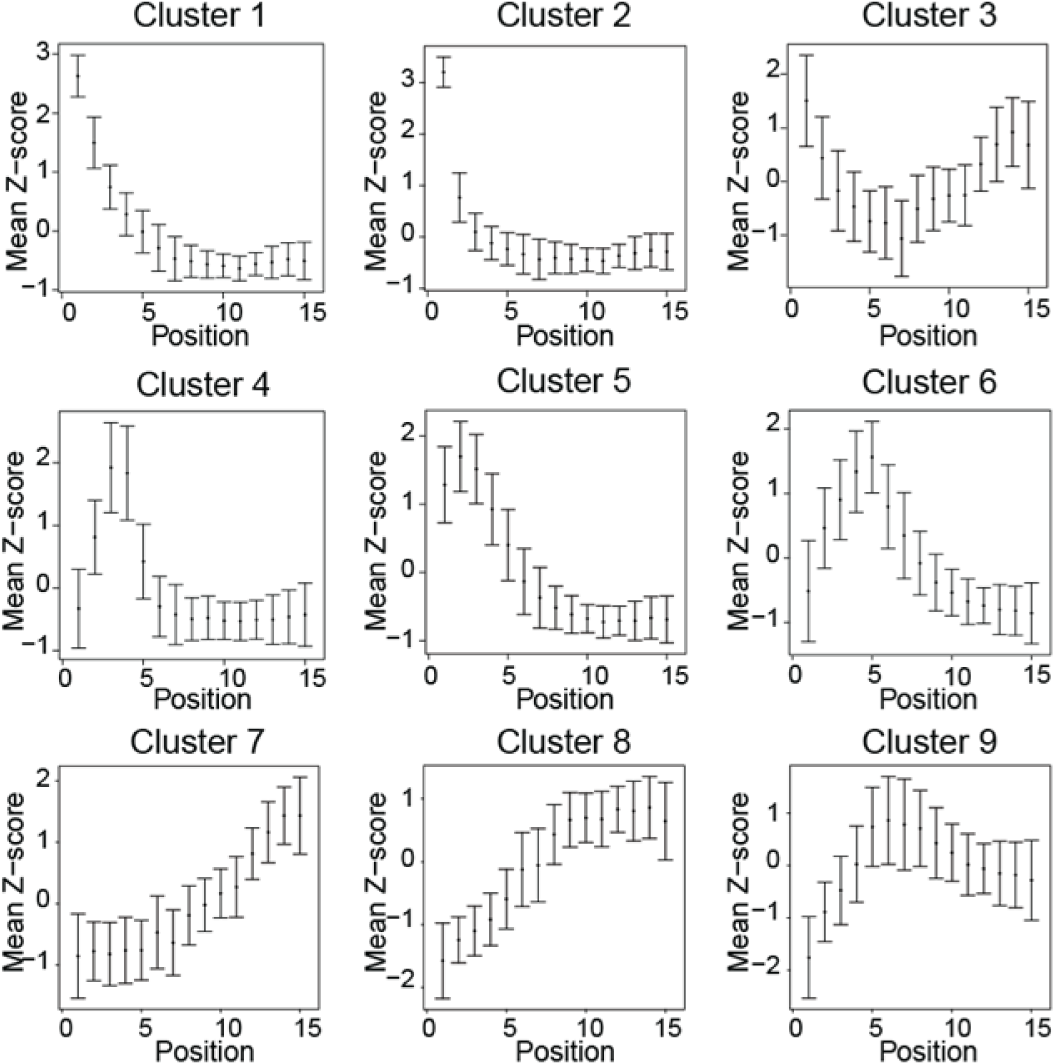
Expression dynamics representing nine developmental regulons during photosynthetic differentiation. RNA-seq data from 15 sections along the length of the maize seedling leaf blade (Wang et al., 2014) was *z-*scored across the leaf gradient, and the *z-*scores were used for *k-* means clustering (*k*= 9). The mean and standard deviation of the *z*-scores for all genes in each cluster are displayed from the leaf base (Position 1) to tip (Position 15).

